# A human developmental and adult brain atlas benchmarks dopaminergic stem cell models and cell therapy candidates

**DOI:** 10.64898/2026.06.19.733041

**Authors:** Vittoria D. Bocchi, Ken To, Leslie Weber, Paul Zumbo, Lisa Kim, Iwasaki Yuko Shida, Donghe Yang, Petter Storm, Alessandro Fiorenzano, Edoardo Sozzi, James Hackland, Chuan Xu, Tae Wan Kim, Fani Memi, Nicola J. Drummond, Nadine Bestard-Cuche, Andrea Corsinotti, Omer Ali Bayraktar, Niranjan Sawarkar, Regine S. Tipon, Kavitha Krishna Sudhakar, Aaron Zhong, So Yeon Koo, Jinghua Piao, Xiaoling He, David Horsfall, Daniela Basurto-Lozada, Ting Zhou, Viviane Tabar, Joseph E. Powell, Roger A. Barker, Barbara Treutlein, Gist Croft, Asuka Morizane, Tilo Kunath, Malin Parmar, Sandra Blaess, Rajeshwar Awatramani, Doron Betel, Sarah A. Teichmann, Lorenz Studer

## Abstract

Parkinson’s disease (PD) is characterized by the progressive loss of midbrain dopaminergic (mDA) neurons^1^. Stem cell–derived mDA neurons hold promise for disease modelling^2,3^ and are currently in clinical trials for cell replacement therapy^4–6^. However, systematic benchmarking has been limited by the lack of a unified high-resolution reference and methods that quantify incomplete or mixed lineage specification *in vitro*^7^. We establish a single-cell and spatial atlas of the human developing diencephalon–midbrain–hindbrain axis resolving 93 cell subtypes, including 39 lacking prior single-cell characterization and 4 entirely novel populations. Using this atlas as a reference, we integrate 19 hPSC-derived mDA datasets, both published^2,8–25^ and unpublished, to build the Human Dopaminergic Neural Atlas (HDNA) spanning 2D, 3D, and graft models, including those used in clinical trials. To classify cells and quantify lineage fidelity, we develop CapybaraBrain, a marker-driven non-negative decomposition framework that assigns each cell continuous identity scores across all 93 developmental programs, enabling systematic discrimination of discrete, transitioning, and cross-lineage hybrid states^26^. We uncover a pervasive landscape of off-target populations reflecting relaxed transcriptional boundaries *in vitro,* including a previously unrecognized TH–PITX2 midbrain neuronal population, and we validate atlas-predicted latent lineage plasticity through inducible genetic fate mapping in mouse models. We further define maturation-associated transcriptional programs by harmonizing adult mDA subtype atlases, revealing that dopaminergic identity and maturation are partially decoupled across protocols. Finally, projecting PD patient-derived tri-cultures onto the HDNA uncovers genotype- and cell-type-specific transcriptional dysregulation. Together, these integrated atlases and computational framework establish a unified standard for benchmarking differentiation fidelity, exposing off-target states, and guiding next-generation PD models and cell therapies.

## MAIN

PD is characterized by the progressive degeneration of midbrain dopaminergic (mDA) neurons in the substantia nigra^1^ that leads to motor symptoms such as bradykinesia, tremor, and rigidity^27^. This has driven efforts to generate human mDA neurons from pluripotent stem cells for disease modelling^2,3^ and cell replacement therapy^4–6^. For disease modelling, a growing range of systems is now used including monolayer cultures^2,8–11^, organoids^12,13,15,28,29^ and assembloids^18,20^. In parallel, transplantation strategies have focused on protocols that generate floor plate progenitors for grafting^11,14,17,23^. As several stem cell–based therapies are now in the clinic^6,30–33^, together with the increasing use of these systems for mechanistic studies, compound screening, and neurotoxicity testing^34–36^, it is imperative to define the advantages, limitations, and optimal applications of each protocol and model.

Single-cell profiling enables detailed characterization of cell subtypes and key differences between protocols in generating mDA neurons. Although single-cell profiling has increasingly been applied to stem cell-derived dopaminergic protocols, most studies remain protocol-specific^2,14,17,21–23,37^, while the few comparative analyses across multiple protocols^38,39^ have relied on broad regional annotations or midbrain-centered reference frameworks with limited resolution of adjacent brain regions. This limitation is particularly important because stem cell-derived dopaminergic cultures can exhibit patterning drift both within the midbrain and into neighboring diencephalic and hindbrain territories^11,21–23^. A clinically relevant example of why precise subtype classification is important is the potential emergence of neighboring hindbrain serotonergic neurons through patterning drift. These off-target cells were detected in fetal ventral mesencephalic grafts used in PD transplantation trials^40–42^ and have been implicated in graft-induced dyskinesias in a subset of patients^41^. However, serotonergic neurons are unlikely to be the only relevant off-target population *in vitro*, as multiple neighboring neuronal identities may arise during differentiation with currently unknown functional consequences. This illustrates how accurately resolving subtype identity is essential for assessing the fidelity and safety of stem cell-derived cell products.

The reference atlases currently available to benchmark these protocols remain limited in several respects. First, existing datasets often apply inconsistent nomenclature to define midbrain subtypes^43,44^, complicating direct comparison across studies. Second, neighboring diencephalic and hindbrain regions are frequently represented by broad annotations^44^ that do not resolve the underlying dorsoventral and rostro caudal diversity of progenitor and neuronal subtypes. Although single-cell datasets of selected diencephalic regions are available^45,46^, these remain fragmented across studies, whereas the hindbrain, outside of the cerebellum^47,48^, has been under-characterized at single-cell resolution despite harboring substantial cellular diversity^49–51^. Together, these limitations make it difficult to determine the precise identity of recurrent off-target populations generated in stem cell-derived dopaminergic products. Beyond resolving off-target identities, there is also a critical need to assess the maturation state of the dopaminergic neurons themselves. Without a unified framework to quantify maturation across models, it is challenging to determine whether differences across protocols reflect improved specification, enhanced maturation, or both. This challenge is hindered by current adult post-mortem human dopaminergic neuron datasets^52,53^, which profile individuals of different ages and often use differing subtype nomenclatures and annotation schemes, hindering clear and consistent assessment of maturation across studies.

A deeper conceptual challenge is that stem cell-derived models often exhibit incomplete lineage specification *in vitro*^7,23^, requiring tailored computational approaches capable of resolving cells with hybrid transcriptional profiles^26,54^. Current large atlas efforts of stem cell-derived products^12,55^ have largely overlooked this issue by imposing discrete labels onto inherently continuous/mixed cellular states, thereby obscuring mixed transcriptional programs and limiting the detection of lineage drift and hybrid cell states across models. Addressing these challenges requires integrated developmental and adult consensus atlases, coupled with a unified *in vitro*-derived atlas and dedicated computational tools, to resolve continuous and hybrid identities, benchmark stem cell-derived dopaminergic models, and guide future refinement for PD research and cell therapy.

## RESULTS

### A unified single-cell and spatial atlas of the human developing diencephalon–midbrain–hindbrain axis reveals novel and under-resolved cell subtypes

We assembled a comprehensive first- and second-trimester human developing caudal brain atlas by integrating diencephalon, midbrain and hindbrain datasets from the Human Cell Atlas (HCA) Development v1.0^18^ with ∼60,000 additional developing midbrain and hindbrain cells (Fig. 1a, Supplementary Table 1). In the midbrain, we resolved cross-study annotations^43,44^ inconsistencies using CellHint^56^ to establish consensus cell labels and refine fine-grained subtypes (Extended Data Fig. 1a,b). We next resolved regional annotations across the diencephalon and hindbrain, transitioning from the broad cell-subtype framework of HCA v1 to a high-resolution subtype atlas (Fig. 1a-d, Extended Data Fig. 2a-d). In the diencephalon, this refinement resolved thalamic, prethalamic, subthalamic, hypothalamic, habenula, and zona incerta lineages (Extended Data Fig.2 a-b). In the hindbrain, we uncover extensive molecular diversity not previously resolved at single cell resolution, defining serotonergic, noradrenergic, motor and interneuron populations, the Bötzinger complex, pre cerebellar nuclei, and cerebellar lineages from the ventricular zone and rhombic lip, together with diverse V2 populations that include entirely novel subsets (Fig.1 a-d, Extended Data Fig. 2c–d).

**Figure 1.**
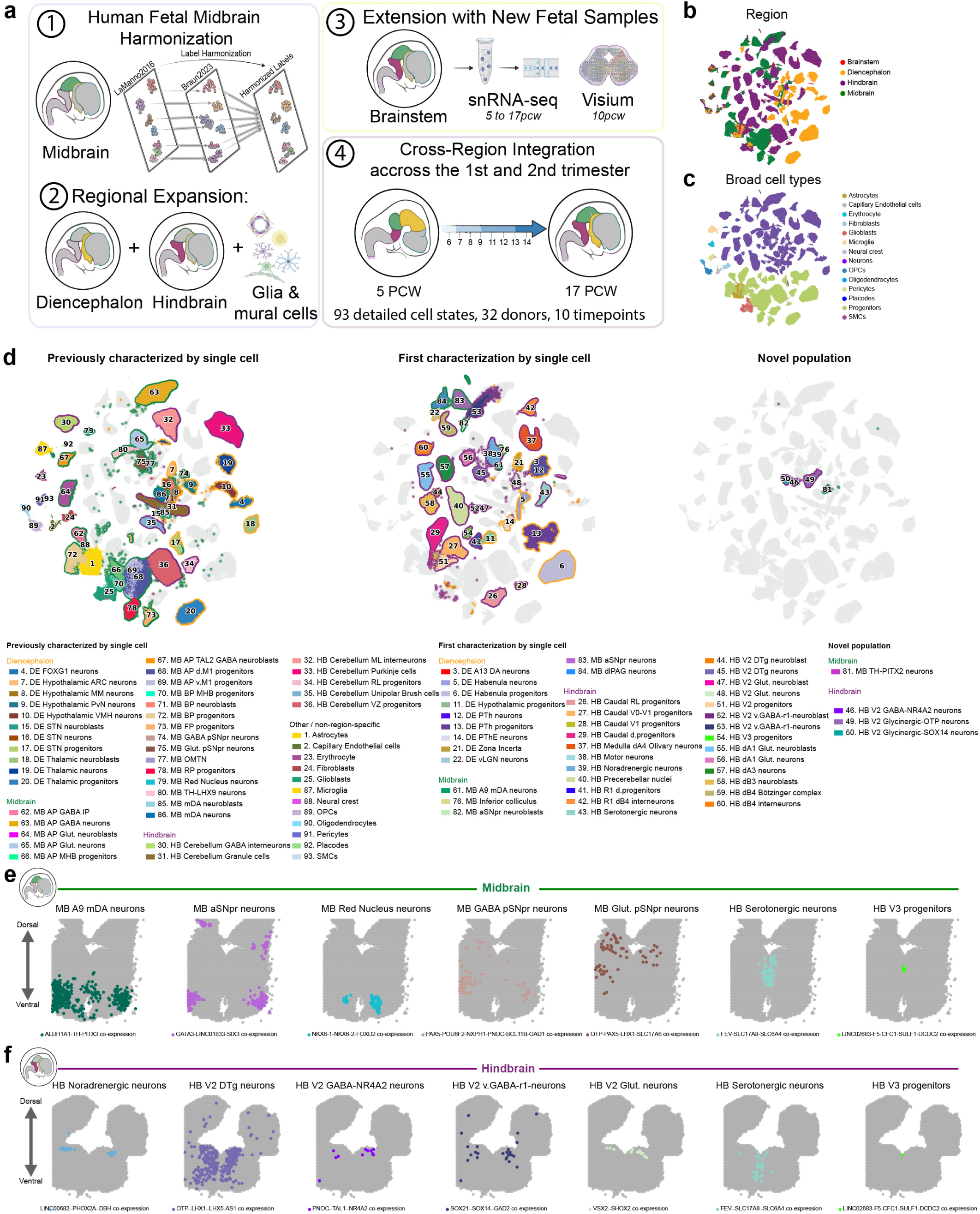
Integrated single-cell and spatial atlas of the human fetal caudal brain. **a**, Schematic overview of the workflow used to construct the human fetal brain atlas. **b-c**, UMAP visualization coloured by dissection region (b) and annotations from the Human Cell Atlas Development fetal brain collection version 1 with broad cell labels (c). **d**, UMAP of fetal cell states grouped by prior level of characterization: previously described, newly fully characterized, and novel populations. Coloured halos indicate regional origin (diencephalon, midbrain, hindbrain), while non–region-specific cell types have no halo. **e-f**, Spatial co-expression of cell-defining marker genes across voxels, illustrating the spatial distribution of specific subtypes in the midbrain (e) and hindbrain (f) in 10 PCW sections.

**Figure 2.**
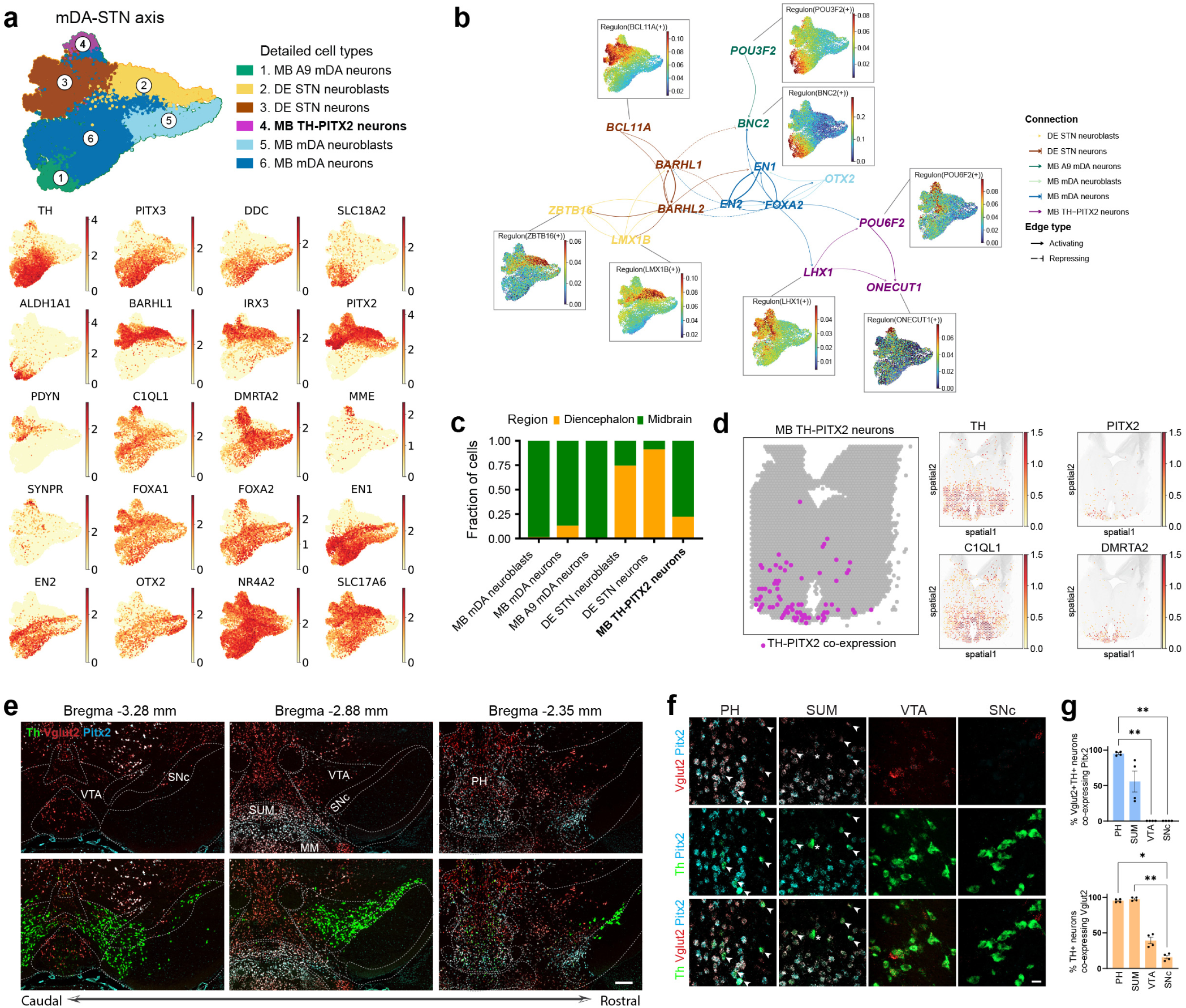
Gene regulatory network and spatial profiling of a novel TH–PITX2 neuronal population. **a,** Subclustering of mDA–STN neurons identifies a TH–PITX2 neuronal population with an intermediate transcriptional profile, shown by canonical and novel mDA (*TH*, *PITX3*, *DDC*, *SLC18A2*, *ALDH1A1*) and STN (*BARHL1*, *IRX3*, *PITX2*, *PDYN*, *C1QL1*, *DMRTA2*, *MME*, *SYNPR*) together with canonical midbrain floor plate (*FOXA1*, *FOXA2*, *EN1*, *EN2*, *OTX2*, *NR4A2*, *SLC17A6*) markers. **b**, SCENIC gene regulatory networks highlighting novel transcription factors (TFs) in mDA, STN, and TH–PITX2 neurons. Nodes represent key TFs, and edges indicate predicted regulatory interactions. Edge colors correspond to cell-type specificity. Inset UMAPs show the activity of representative regulons for each subtype. **c,** Stacked bar plot showing regional distribution of STN, mDA and TH–PITX2 neurons in different regions. **d**, Spatial validation of TH–PITX2 neurons in a 10 PCW fetal midbrain section showing spatial co-expression of specific markers (left panel) and individual gene expression spatial profiles across voxels (right panel). **e**, RNAscope in situ hybridization of Pitx2, Vglut2 (Slc17a6), and Th overlaid onto coronal sections from the Allen Mouse Brain Atlas. Scale bar: 200 μm. **f**, High magnification images of RNAscope in situ hybridization. Pitx2 is expressed in Vglut2+Th+ (arrowheads) and Vglut2-Th+ neurons (asterisk). Scale bar: 20 μm. **g**, Percentage of Vglut2+Th+ neurons co-expressing Pitx2 and Th+ neurons co-expressing Vglut2 in the PH, SUM, VTA, and SNc (Kruskal-Wallis with Dunn’s test). Significant differences at *p < 0.05, **p < 0.01, and ****p < 0.0001. n (sections; mice) = 18; 4. Data are presented as mean ± SEM. Abbreviations: PH, posterior hypothalamic nucleus; SUM, supramammillary nucleus; SNc, substantia nigra pars compacta; VTA, ventral tegmental area.

This iterative integration of new data (Extended Data Fig. 3a–f) and refinement yielded a high-resolution atlas comprising 896,761 cells spanning 93 cell subtypes across the developing human caudal brain. Of these, 50 correspond to previously characterized single-cell populations in different scattered studies, whereas 39 represent cell subtypes that lacked prior single-cell resolution in the human developing brain and 4 constitute previously unreported populations (Fig. 1d), including a novel TH-PITX2 neuronal subtype. Given the overlap of subtype markers in the literature (Extended Data Fig. 1b, 2b,d), we derived a dictionary of highly specific marker genes for each cell subtype (Extended Data Fig.4 a-d, Supplementary Table 2) to enable precise discrimination between closely related identities. We next mapped these subtypes into their anatomical context using spatial transcriptomics, revealing the organization of both canonical and previously under-resolved population. In the midbrain, we identified refined molecular signatures of mDA neurons, including co-enrichment of *TH* and *PITX3* with the previously unknown markers *LINC00616* (Extended Data Fig. 5a), and resolved their spatial redistribution by 10 post-conception weeks (PCW) toward lateral ventral domains (Fig. 1e, Extended Data Fig. 5 b,c).

**Figure 3.**
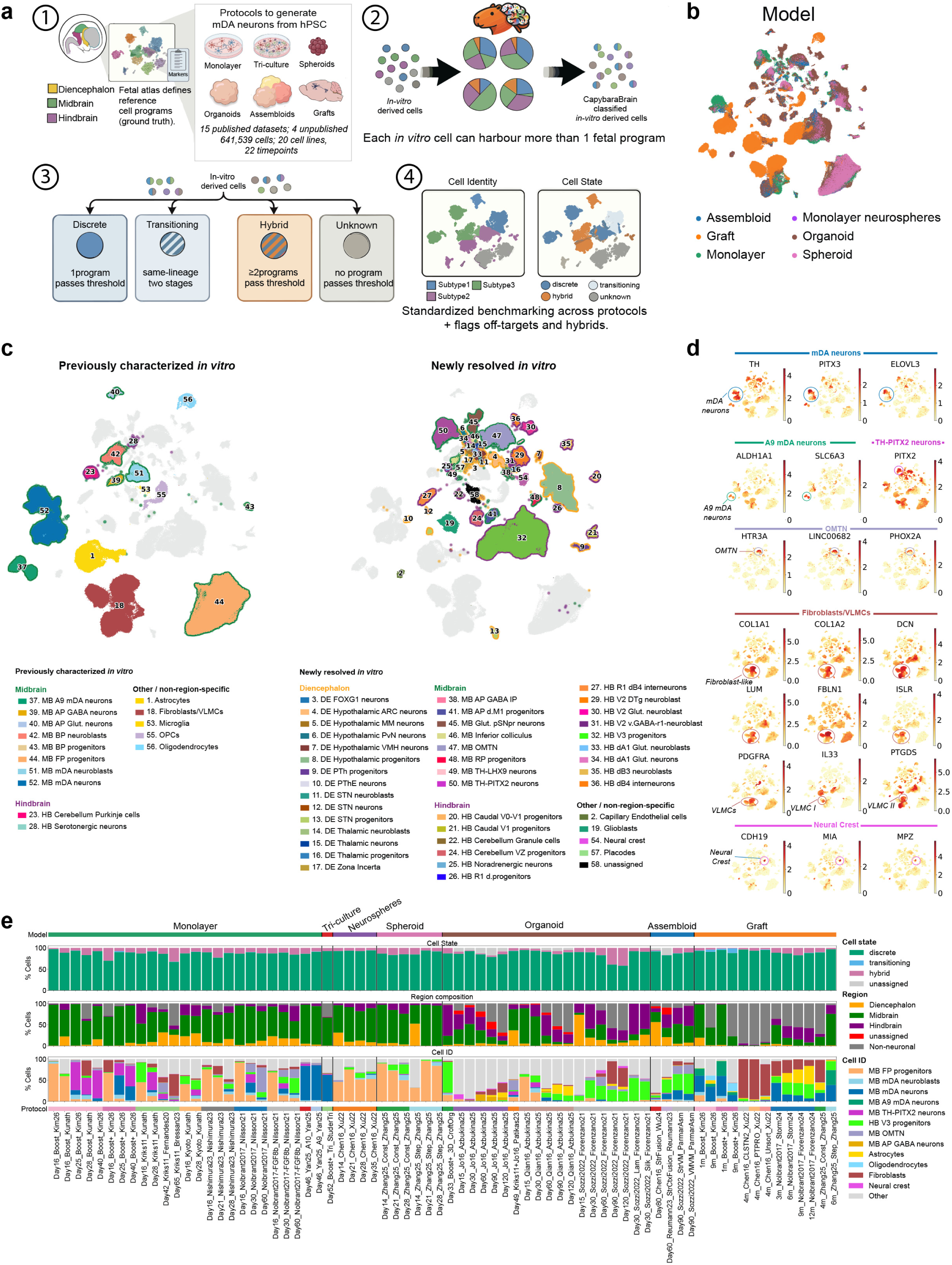
Construction and classification of the Human Dopaminergic Neural Atlas (HDNA) across stem-cell derived mDA differentiation protocols and models. **a,** Schematic overview of the CapybaraBrain workflow, illustrating reference-based decomposition of *in vitro* profiles, cell subtype and state classification, and majority-voting refinement. **b,** UMAP of all integrated *in vitro* datasets coloured by model type. **c**, UMAP visualization of in vitro cells coloured by CapybaraBrain cell type annotation. Left panel: previously characterized cell types in vitro. Right panel: previously unresolved (novel) cell types identified in this study. Coloured halos indicate regional origin (diencephalon, midbrain, hindbrain), while non–region-specific cell types have no halo. **d,** UMAP showing expression of celltype specific markers for canonical mDA neurons, A9 mDA neurons, TH–PITX2 neurons, OMTN, fibroblasts, and neural crest cells. **e,** Stacked bar plots summarizing CapybaraBrain classifications across protocols and timepoints. Top panel shows cell state distribution; middle panel shows inferred regional origin; bottom panel shows major cell identities defined by majority voting. Each bar represents one Timepoint–Protocol–Publication combination, grouped by model type (coloured band). Percentages indicate the proportion of cells assigned to each category within that dataset.

**Figure 4.**
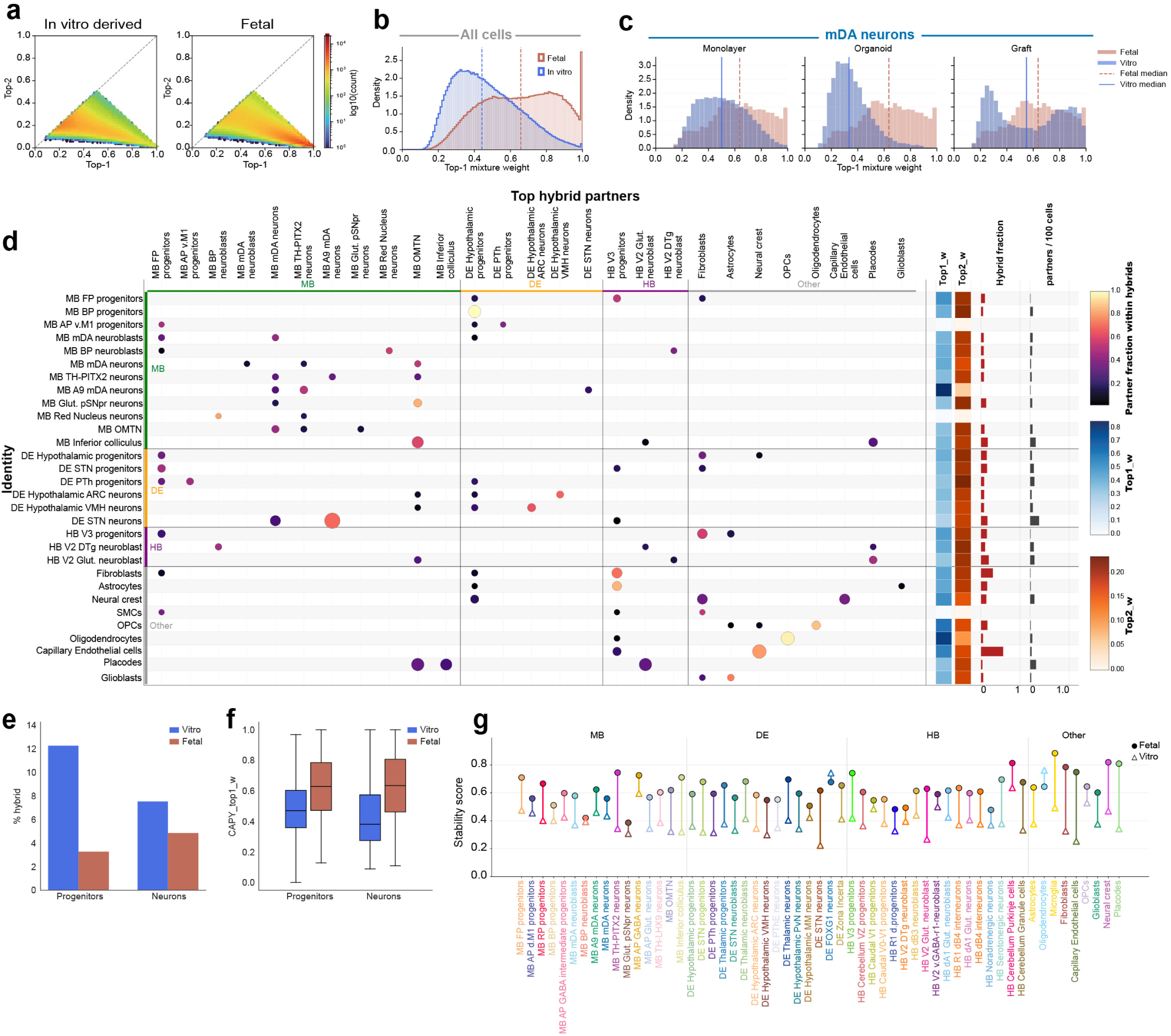
CapybaraBrain-based quantification of lineage dominance and hybridization *in vitro* and in fetal cells. **a**, Hexbin density plot of Top-1 versus Top-2 CapybaraBrain mixture weights for *in vitro*-derived and fetal cells (log10 scale). The dashed diagonal indicates equal contribution of primary and secondary lineage programs. **b**, Histogram of Top-1 CapybaraBrain mixture weights for fetal and *in vitro*–derived cells. Vertical lines indicate median Top-1 weights. **c**, Distributions of Top-1 CapybaraBrain mixture weights for MB mDA neurons and MB A9 mDA neurons across *in vitro* model systems, shown alongside the fetal reference in each panel. Vertical lines indicate median Top-1 weights. **d,** Dot plot showing the most frequent hybrid partners for each cell type. Each row represents a cell type, and each column shows its most common hybrid partners. Each dot indicates a hybrid combination between two cell types. Dot size reflects how often this hybrid occurs, specifically the percentage of cells of a given type that form this hybrid. Dot color shows how strongly a partner contributes within the hybrid, with darker colors indicating a stronger contribution. The coloured side bars group cell types by brain region (midbrain, diencephalon, hindbrain). The panels on the right summarize additional features for each cell type: the average strength of the primary (Top1) and secondary (Top2) programs, the fraction of cells that are hybrids, and the number of different hybrid partners. Only cell types with sufficient representation in hybrid states are shown (≥15 cells per identity and ≥20 cells per hybrid pair). For each cell type, only the top three hybrid partners are displayed, defined as those with the highest fractional contribution within hybrid states. **e**, Bar plot showing the percentage of hybrid cells within progenitor and neuronal populations in fetal and in vitro datasets. **f**, Box plots showing the distribution of Top1 identity weights (CAPY_top1_w) for progenitor and neuronal populations in fetal and in vitro datasets. **g**, Dumbbells showing the stability of each cell state, defined as mean Top1 identity weight × (1 − hybrid fraction), where higher values indicate stronger, more consistent identity and lower hybridization. Each vertical dumbbell connects matched cell states between fetal (filled circles) and in vitro (open triangles) datasets.

Spatial mapping indicates that by 10 PCW, diencephalic-derived SIX3⁺ anterior substantia nigra pars reticulata neurons (aSNpr)^57^ marked by the novel marker *LINC01833* have reached their final positions, whereas hindbrain-derived glutamatergic and GABAergic posterior SNpr (pSNpr) populations defined by *OTP*, *LHX1*, *BCL11B* and *PNOC* are migratory at this stage (Fig. 1e, Extended Data Fig. 5 b,c). Finally, we identified serotonergic neurons and progenitors along the ventral midline at 10 PCW and in close proximity to mDA neurons (Fig. 1e, Extended Data Fig. 5 b,c). These cells are consistent with the rostral serotonergic lineage that gives rise to the dorsal raphe nucleus^58,59^. The presence of serotonergic progenitors within our caudal midbrain section suggests the presence of a wave of serotonergic neurogenesis along the midbrain ventral midline, rather than its confinement to the hindbrain as seen in mouse studies^60^. This shows that serotonergic neurons reported in fetal ventral mesencephalic grafts used in PD clinical transplantation trials^40–42^, that cause graft-induced dyskinesias^41^, are derived from the caudal midbrain areas.

In the hindbrain, spatial validation confirmed both established and newly resolved populations. In the dorsal hindbrain, cerebellar Purkinje neurons expressing *SKOR2*^61^ and *CA8*^62^ confirmed correct r1 localization (Fig. 1f, Extended Data Fig. 5 d,e), while noradrenergic neurons lateral to the fourth ventricle were defined by *PHOX2A* and *DBH*, with *LINC00682* as an additional novel marker (Fig. 1f, Extended Data Fig. 5 d,e). We resolved V2 derived neurons, previously described only in murine models^57^, by co-expression of *OTP*, *LHX1*, *PAX3* and *FOXO1*, and delineated the spatial location of multiple novel V2-derived sub-populations, including GABAergic *NR4A2* neurons (*TAL1*, *NR4A2*) and a ventral r1 GABAergic population (*EN1*, *SOX14*, *OTX1*, *GAD2*, *SOX21*), alongside glutamatergic V2 neurons defined by *VSX2* and *SHOX2* (Fig. 1f, Extended Data Fig. 5 d,e).

Together, this consensus developmental atlas defines a spatially resolved framework of the developing human caudal brain, capturing both established and previously unresolved cell subtypes, and provides a foundation for studying human neurodevelopment, disease vulnerability, and benchmarking stem cell–derived models.

### A novel TH–PITX2 neuronal population is conserved from human development to the adult mouse brain

Given that clinical-grade differentiation protocols^25,63^ are carefully fine-tuned to bias specification to mDA over subthalamic nucleus (STN) specification^64^, and that the novel midbrain TH–PITX2 population identified, occupies an intermediate transcriptional state between these lineages (Fig. 2a), we focused on this population to resolve its molecular program. We found that this population is specifically enriched for *TH*, *PITX2*, *C1QL1*, *DMRTA2*, *MME* and *SYNPR*, while all three lineages share expression of *NR4A2* and the glutamatergic marker VGluT2 (*SLC17A6*) (Fig. 2a). We then used SCENIC^65^ to define key transcription factors (TFs) regulating these three lineages (Fig. 2b). The predicted core regulatory network of mDA neurons is centered on the known transcription factors *EN1*, *EN2*, and *FOXA2*. A9 mDA neurons show selective expression of the novel TFs *BNC2* and *POU3F2* (Fig. 2b). The core mDA FOXA2 module also connects and activates the TH-PITX2 specific transcription factors *POU6F2, LHX1*, and *ONECUT1* (Fig. 2b), suggesting that this population could be midbrain floor plate derived like mDA neurons. This is further reinforced by its location in the midbrain, with a small fraction spanning into the diencephalon (Fig. 2c), likely reflecting minor variability in dissection boundaries across samples or a rostral enrichment of this population near the midbrain/diencephalic border. Spatial studies confirmed that TH-PITX2 neurons are located to the most ventral region of the midbrain at 10 PCW (Fig. 2d).

We next searched for the corresponding population in the adult mouse midbrain^66^, and identified a cluster of cells co-expressing *Th*, *Pitx2, vGlut2, Nr4a1, Lhx1* and *Pou6f2* (Extended Data Fig. 6a). We validated this finding in rostral coronal section of the mouse midbrains, which confirm a population of Th⁺/Pitx2⁺/VGlut2⁺ neurons (Fig.2e). These cells were localized within the posterior hypothalamic region (PH) and the supramammillary nucleus (SUM). Consistent with this localization, quantitative analysis revealed that the majority of Th⁺ neurons in PH and SUM co-express VGlut2 and Pitx2 (Fig. 2f, g), whereas this population was largely absent from the ventral tegmental area (VTA) and substantia nigra pars compacta (SNc). These findings indicate that the newly identified Th⁺-Pitx2⁺-vGlut2⁺ neurons represent a distinct population localized at the rostral midbrain–diencephalic border.

### An *in vitro* multi-model single-cell atlas and cell state deconvolution framework reveal pervasive off-target identities in hPSC-derived dopaminergic neurons

To systematically benchmark the fidelity of stem cell–derived midbrain dopaminergic (mDA) neurons against their *in vivo* counterparts, we constructed the *in vitro* Human Midbrain Dopaminergic Neural Atlas (HDNA) by integrating datasets across monolayer cultures, triculture systems, organoids, assembloids, and grafts, drawn from 15 published and 4 unpublished studies, including all the protocols used in human clinical trials (Extended Data Fig. 7a, Supplementary Table 3). Following quality control, the integrated atlas comprised 641,539 cells with 20 different cell lines (both ESCs and iPSCs) and 22 unique timepoints (Extended Data Fig. 7 a,b). To classify cells and quantify incomplete lineage specification in vitro^7,23^, we developed CapybaraBrain (see Methods), a marker-driven (Supplementary Table 4) non-negative least squares (NNLS) framework that decomposes each cell’s transcriptional profile into weighted contributions from all 93 developmental reference programs. This enables three capabilities unavailable to conventional single-label classifiers: quantitative measurement of program fidelity per cell, detection of off-target identities, and, critically, discrimination between lineage-consistent transitioning states and cross-lineage hybrid mixtures (Fig. 3a). Benchmarking against CellTypist^67^ classification on human developmental data, showed superior classification accuracy (Extended Data Fig. 7c), validating the framework before application to the HDNA.

Across the HDNA, CapybaraBrain identified 58 distinct cell subtypes, including 15 previously described in hPSC-derived dopaminergic systems and 43 not previously resolved *in vitro* (Fig. 3c). To enable cross-dataset comparison at this resolution, we performed CapybaraBrain label-aware integration using CellHint^56^, which outperformed other methods (Extended Data Fig. 7d). Across neuronal populations, canonical mDA neurons defined by high expression of *TH*, *PITX3* and *ELOVL3* (Fig. 3d) were detected predominantly in monolayer protocols and grafts, but were reduced in 3D systems (Fig. 3e). A more mature subset of A9 mDA neurons marked by *ALDH1A1* and the dopamine transporter DAT (*SLC6A3*) was observed exclusively in grafted cells (Fig. 3e), indicating that only the *in vivo* environment promotes refinement of dopaminergic subtype identity. In addition to intended mDA lineages, we identified multiple previously unrecognized off-target neuronal populations *in vitro*, originating from adjacent developmental territories to the midbrain dopaminergic domain, indicating imprecise regional patterning. In particular, many monolayer protocols preferentially generated the newly identified TH–PITX2 neuronal population (Fig. 3e), characterized by elevated *PITX2* and low *PITX3* expression relative to canonical mDA neurons (Fig. 3d), a finding further supported by immunofluorescence in the Boost+ protocol at day 40, confirming the presence of TH⁺, PITX2⁺ and FOXA2⁺ neurons (Extended Data Fig. 7e). Other off-targets include OMTN and glutamatergic pSNpr identities, alongside smaller fractions of additional neuronal subtypes. Notably, the predominant off-target population corresponded to hindbrain V3 progenitors (Fig. 3c,e), a neighbouring progenitor to the intended midbrain floor plate progenitor. These cells were consistently observed across protocols and timepoints, indicating that many differentiation strategies promote caudalization toward ventral hindbrain identities rather than maintaining midbrain-specific patterning.

Beyond neuronal populations, diverse non-neuronal and off-target lineages were detected. Fibroblast-like cells expressing *COL1A1*, *COL1A2*, *DCN* and *LUM* were prevalent, particularly in graft-derived datasets, but also emerged in monolayer and organoid conditions at later stages. In grafts, these cells closely resemble the fibroblast-like cell in the brain defined as vascular leptomeningeal cells (VLMCs) described previously^14,17,22,37^, marked by *PDGFRA* expression. Notably, we further resolved two distinct VLMC subtypes consistent with those reported in the mouse brain^68^, characterized by expression of *IL33* and *PTGDS*, respectively. Surprisingly, neural crest-like populations marked by *CDH19*, *MIA* and *MPZ* were also observed, indicating additional lineage divergence. Astrocytes arose predominantly at later stages and were enriched in organoids, assembloids and grafts, consistent with delayed glial maturation^69–71^, whereas oligodendrocytes were restricted to a subset of grafts^37^. Overall, this exposes a major and previously unrecognized source of mis-specification that is only detectable through this high-resolution developmental reference.

### Incomplete lineage resolution *in vitro* results in hybrid identities through partial engagement of off-target developmental programs

To quantitatively assess lineage fidelity, we compared CapybaraBrain weights, defined as the proportional contribution of each reference cell state program to a given cell’s transcriptional profile, between *in vivo* and *in vitro*-derived datasets. Fetal cells exhibited a stronger enrichment toward high Top-1 values (the highest contributing reference program), indicating dominant assignment to a single cell program. In contrast, *in vitro* cells were shifted toward lower Top-1 weights with correspondingly higher contributions from secondary programs (Top-2), consistent with more mixed lineage identities (Fig. 4a,b; Extended Data Fig. 7f). Accordingly, we observe a higher proportion of discrete identities in *in vivo*, whereas *in vitro* datasets exhibited a higher number of hybrid states (Extended Data Fig. 7g). We did not observe strong differences in discrete versus hybrid states between models (Extended Data Fig. 7h), suggesting that increased lineage mixing is a general feature of most models. However, when focusing specifically on Top-1 weights of mDA neurons, graft-derived cells exhibited the highest mDA program dominance, whereas organoid-derived neurons showed the lowest (Fig 4c). This suggests that grafts can achieve stronger lineage commitment, possibly due to *in vivo* environmental cues that promote more complete lineage specification.

Global *in vitro* hybridization topology reveals a structured, regionally constrained, and non random logic, reflecting partial engagement of developmental programs from anatomically and developmentally related subtypes (Fig. 4d). Hybrid relationships are enriched within regional domains, most prominently within the ventral midbrain (MB), but extend across ventral diencephalic and hindbrain boundaries, particularly in progenitor populations, revealing lineage coupling that can be quantitatively captured and compared. For instance, MB floor plate progenitors hybridize toward hindbrain V3 and hypothalamic progenitor programs, reflecting erosion of anterior posterior identity constraints. Conversely, HB V3 progenitors show persistent coupling to MB floor plate progenitors, consistent with instability at the midbrain hindbrain boundary (Fig. 4d). Among neuronal populations, mDA neurons exhibit contaminating signatures from neighboring rostral TH–PITX2 neurons and OMTN lineages (Fig. 4d), suggesting that progenitor level instability propagates into neuronal populations, resulting in mixed identities along the rostro caudal axis. Notably, specific ventral progenitor populations, including STN, hypothalamic, midbrain floor plate, and HB V3 progenitors, display strong and recurrent hybridization with non neuronal identities, indicating systematic engagement of glial and mesenchymal transcriptional programs. This pattern is not observed in progenitors occupying more dorsal domains, such as prethalamic and alar plate populations, nor in later basal plate progenitors, indicating a domain specific lineage coupling selectively associated with the ventral developmental contexts. The frequent and directional nature of these hybrid relationships, together with lower identity dominance, shows that *in vitro* cells remain partially specified rather than reaching stable, final identities. Consistently, hybrid states are globally increased *in vitro*, accompanied by reduced Top-1 weights across both progenitor and neuronal populations (Fig. 4e,f), demonstrating persistent lineage mixing throughout differentiation. In line with this, stability analysis across subtypes (Fig. 4g) shows a widespread reduction in identity resolution *in vitro* compared to *in vivo* in most subtypes. However, this effect is not uniform with mDA neurons displaying higher stability, indicating more robust specification, whereas alternative off-target lineages show pronounced instability. Collectively, this defines a “regulatory syntax” of *in vitro* identity, in which cells combine partial elements of multiple lineage programs in a constrained manner, capturing the permissible transitions encoded in the developmental atlas and providing a quantitative framework to measure how far *in vitro* systems deviate from *in vivo* reference states.

### Hybrid lineage coupling reveals latent astrocytic and mesenchymal plasticity in ventral progenitors *in vitro* and *in vivo*

We hypothesized that the hybrid states observed *in vitro* could reflect either an *in vitro* artefact or early and permissive lineage configurations that are normally transient or tightly constrained during development but become detectable through our high-resolution atlas and quantitative decomposition framework. To explore these patterns, we focused on HB V3 progenitors and MB floor plate progenitors, which showed prominent deviations from *in vivo* identity, including both cross-regional coupling and non-neuronal drift (Fig. 5a,b). Both populations displayed hybridization toward fibroblast signatures, with V3 progenitors additionally exhibiting coupling to astrocytic programs. Notably, these non-neuronal signatures were largely absent at the developmental stages analysed, suggesting that their activation *in vitro* reflects either aberrant lineage plasticity or the sustained engagement of programs that may not be captured in the samples sampled. These fibroblast and astrocytic programs are not uniformly distributed but instead show clear model- and protocol-specific patterns. Monolayer cultures display a marked enrichment of fibroblast-hybrid states, whereas astrocytic hybrids are preferentially enriched in organoid and graft systems (Fig. 5c). Midbrain FP progenitor–fibroblast hybrids are found mostly in monolayer protocols while HB V3 progenitor–fibroblast hybrids are observed across most protocols and models and increase over time, paralleling expansion of the V3 progenitor population (Fig. 5d-e, Extended Data Fig. 8a, b), in contrast to MB floor plate progenitors, which rapidly decline during differentiation (Fig. 5f). Together, these patterns reveal both context-dependent lineage biases and conserved temporal dynamics across protocols. An exception was observed in the Xu2022^11^ dataset, which showed reduced fibroblast hybridization *in vitro* (Fig. 5d). However, V3–fibroblast hybrids emerged in the 4-month graft alongside an expanded fibroblast-like population (Fig. 5d, Extended Data Fig. 8b), indicating that this potential is retained but temporally delayed. Consistent with this, early transition to floating neurosphere culture may limit adhesion-dependent cues that promote fibroblast-like programs, whereas in 3D systems such transitions occur later (Fig. 5d). Despite this variability, the persistence of fibroblast/VLMC populations in grafts suggests an intrinsic capacity of both V3 and midbrain floor plate progenitors to engage mesenchymal-like programs at later stages.

**Figure 5.**
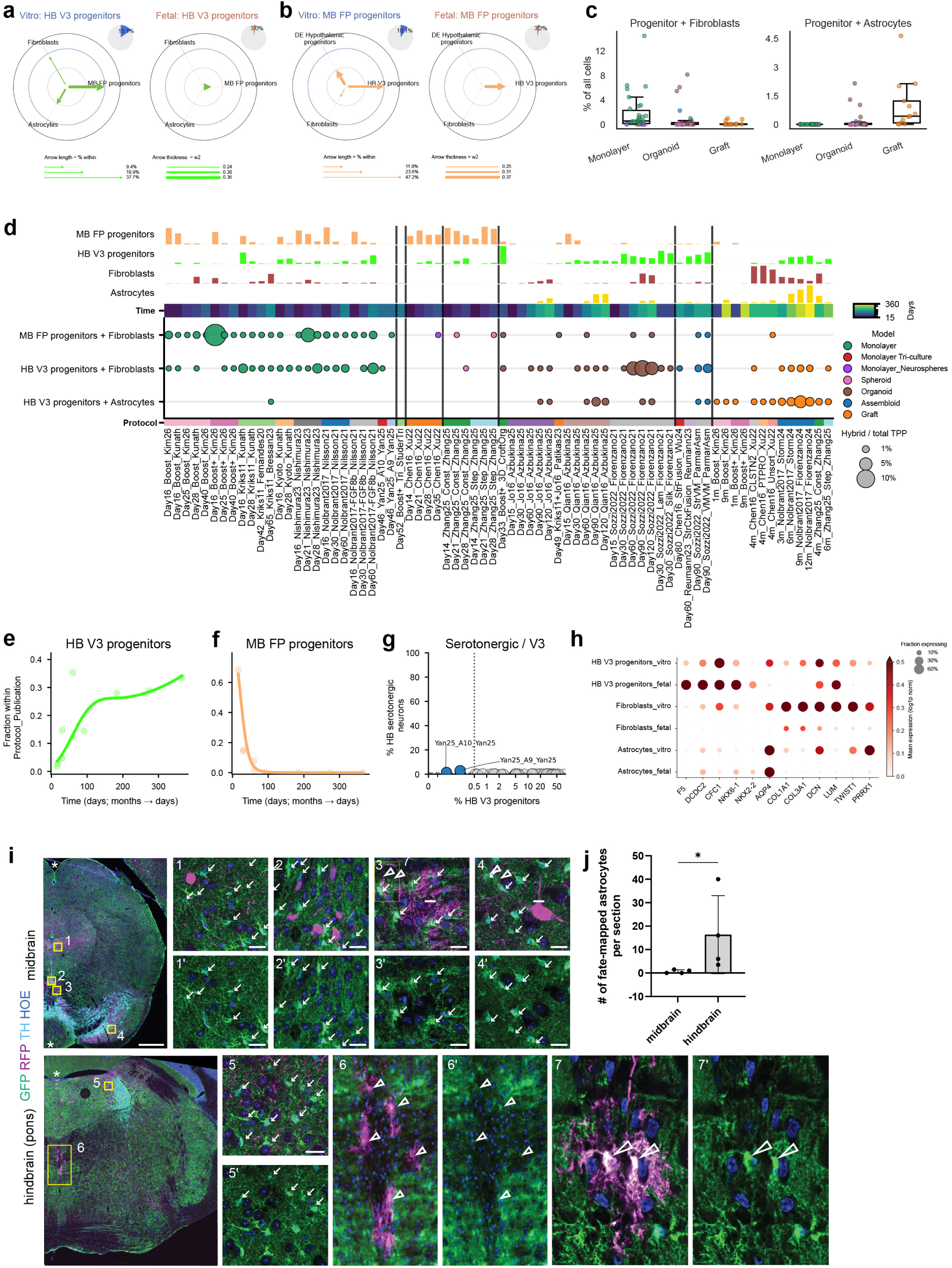
Protocol-dependent hybrid progenitor states associate with fibroblast and astrocyte programs and predict nested lineage potential *in vivo*. **a-b**. Radar plots summarizing the most frequent secondary programs (Top2) associated with hybrid calls for HB V3 progenitors (a) and MB floor plate (FP) progenitors (b), comparing *in vitro* (left) and fetal (right) datasets within each panel. Arrow length is proportional to the percentage of hybrids within the indicated Top1 population (% within), and arrow thickness is proportional to the mean secondary mixture weight (w2), reflecting the strength of the secondary transcriptional contribution. Pie charts represent percentage of hybrids within each progenitor in vitro or in fetal cells. **c**, Boxplots showing the fraction of cells, per timepoint–protocol–publication, classified as hybrids between progenitor programs (hypothalamic progenitors, HB V3 progenitors, and midbrain FP progenitors) and Fibroblasts (left) or Astrocytes (right). **d**, Distribution of progenitor populations and associated hybrid states across differentiation conditions. Top panels show the percentage of selected cell types (midbrain floor plate progenitors, hindbrain V3 progenitors, fibroblasts, and astrocytes) within each condition. The heatmap indicates the corresponding timepoint (in days). The bubble plot shows hybrid relationships between progenitor populations and non-neuronal cell types. Circle size reflects how frequent the hybrid is, calculated as the fraction of all cells in that condition. **e-f**, Fraction of HB V3 (e) and MB FP (f) progenitors across differentiation timepoints within Nolbrant-based protocols. Each dot represents the fraction of cells classified as HB V3 or MB FP progenitors within a given timepoint–protocol–publication group **g**, Relationship between V3 progenitor abundance and serotonergic neurons across differentiation protocols. Each point represents a protocol (Protocol–Publication combination). Only protocols with detectable progenitors (≥0.5% of cells or ≥10 cells) are included. The x-axis is displayed on a symmetric log scale to resolve low-percentage values. Dashed horizontal lines indicate the mean serotonergic neurons across protocols, and vertical dotted lines mark the V3 progenitor detection threshold (0.5%). Protocols above 1% are highlighted and selectively labelled. **h**, Bubble plot comparing gene expression across matched *in vitro* and fetal populations showing expression of V3 (*F5*, *DCDC2*, *CFC1, NKX6.1, NKX2.2*), astrocytes (*AQP4*), fibroblast (*COL1A1*, *COL3A1*, *DCN*, *LUM*) and EMT (*TWIST1*, *PRRX1*) markers. **i**, Shh-GIFM. *Shh^CreER/+^,R26^Ai9/+^,Aldh1l1-GFP* mouse treated with TM at E8.5. Immunostaining for GFP (ALDH1L1+ astrocytes), RFP (fate-mapped cells) and tyrosine hydroxylase (TH, monoaminergic neurons) in the midbrain (upper panels) and hindbrain (bottom panels). HOE: Hoechst. TH labels dopaminergic neurons in the midbrain and noradrenergic neurons in the pons. Asterisks indicate the midline of the sections. (1-7) Higher magnification of the boxed areas in the overview images. In the midbrain RFP+ fate-mapped cells do not overlap with GFP+ astrocytes (arrows); in the hindbrain fate-mapped cells give rise to GFP+ astrocytes along the ventral midline (arrowheads). Scale bars: 500 µm (overview images); 20 µm (higher magnification views in 1-7). **j**, Number of fate-mapped astrocytes in the posterior midbrain and in the anterior hindbrain. 1-3 sections per region; n=3 mice. Error bars: mean ± SD. Mann Whitney test. * p < 0.05.

Given that hindbrain V3 progenitors can give rise to serotonergic neurons *in vivo*^72–74^, an important off-target lineage for cell therapy^41^, we next asked whether these progenitor states are accompanied by their expected neuronal derivatives. Leveraging our atlas, we find that serotonergic neurons are largely absent across differentiation strategies, despite the frequent presence of V3 progenitors (Fig. 5g). Only rare populations of both V3 progenitors and serotonergic neurons were detected in the Yan25 A9 and A10-protocols^21^. These findings suggest that V3 progenitors *in vitro* do not represent progenitors of the serotonergic lineage but instead resemble a closely related and more ventral hindbrain floor plate–like progenitor that in mouse models has been shown to have no neurogenic potential^75^. To investigate this, we examined transcriptional differences between *in vitro* and *in vivo* V3 progenitors and identified shifts in key regulators of neurogenic competence. Compared to *in vivo cells*, *in vitro* V3 progenitors showed absence of *NKX2.2,* an essential transcription factor for serotonergic neuron development^76,77^. In parallel, they display increased expression of astrocytic (*AQP4*) and fibroblast-associated genes (*COL1A1*, *COL1A2*, *DCN* and *LUM*), along with enrichment of EMT-associated transcription factors, including *TWIST1*^78,79^ and *PRRX1*^80^ (Fig. 5h), consistent with activation of mesenchymal-like programs.

To directly test the *in vivo* relevance of lineage potentials predicted by our *in vitro* atlas, we performed genetic inducible fate mapping (GIFM) of Shh floor plate progenitors^81^ at E8.5. We observed that hindbrain floor plate progenitors generate Aldh1l1-positive astrocytes, whereas no astrocytes were detected from corresponding midbrain floor plate progenitors (Fig. 5i,j). These data support our *in vitro*observations that astrocyte generation is restricted to hindbrain V3 floor plate progenitors. Instead, Shh-responsive progenitors (Gli1-GIFM at E9.5)^81^ of both the midbrain and hindbrain label COL1A1 fibroblast-cells surrounding blood vessels specifically in ventral domains (Extended Data Fig. 9a), a finding supportive of the hypothesis that these ventral progenitor populations can give rise to perivasculature fibroblasts. However, we cannot fully exclude the possibility that these perivascular fibroblasts derived from Gli1+ mesenchymal progenitors migrating into this region. Together, these results establish that our *in vitro* atlas captures novel region-specific lineage potentials of ventral progenitors that extend beyond canonical neuronal fates and are supported by *in vivo* genetic fate mapping.

### A unified adult atlas framework reveals decoupled dopaminergic identity and maturation and enables benchmarking of stem cell–derived PD models

To enable systematic assessment of dopaminergic identity and maturation, we generated an integrated atlas of adult mDA neuron subtypes by harmonizing two recent post-mortem studies^52,53^ (Fig. 6a, Extended Data Fig. 10a, Supplementary Table 5) and established a unified nomenclature based on the core genes shared by each subtype (Extended Data Fig. 10b). Using CellHint^56^, we integrated these datasets into a single consensus reference atlas spanning young, adult and elderly samples (Fig. 6b-d, Extended Data Fig. 10c). We next focused on A9 mDA neurons and integrated developmental (6–17 PCW) and young adult (29–50 years) datasets (Fig. 6e), defining stage-specific regulomes and gene modules that distinguish developing from mature states (Fig. 6f,g, Extended Data Fig. 10d, Supplementary Table 6). These findings reveal both conserved regulators previously broadly linked to general neuronal development and maturation (*MBNL2*^82^, *EZH2*^83^, *TCF4*^84^ and *HDAC2*^85^), and previously unrecognized (*ZNF248*, *ZNF483* and *ZNF709*) regulators shaping mature neurons. We then leveraged these signatures to quantitatively assess the maturation state achieved by mDA neurons across the HDNA.

**Figure 6.**
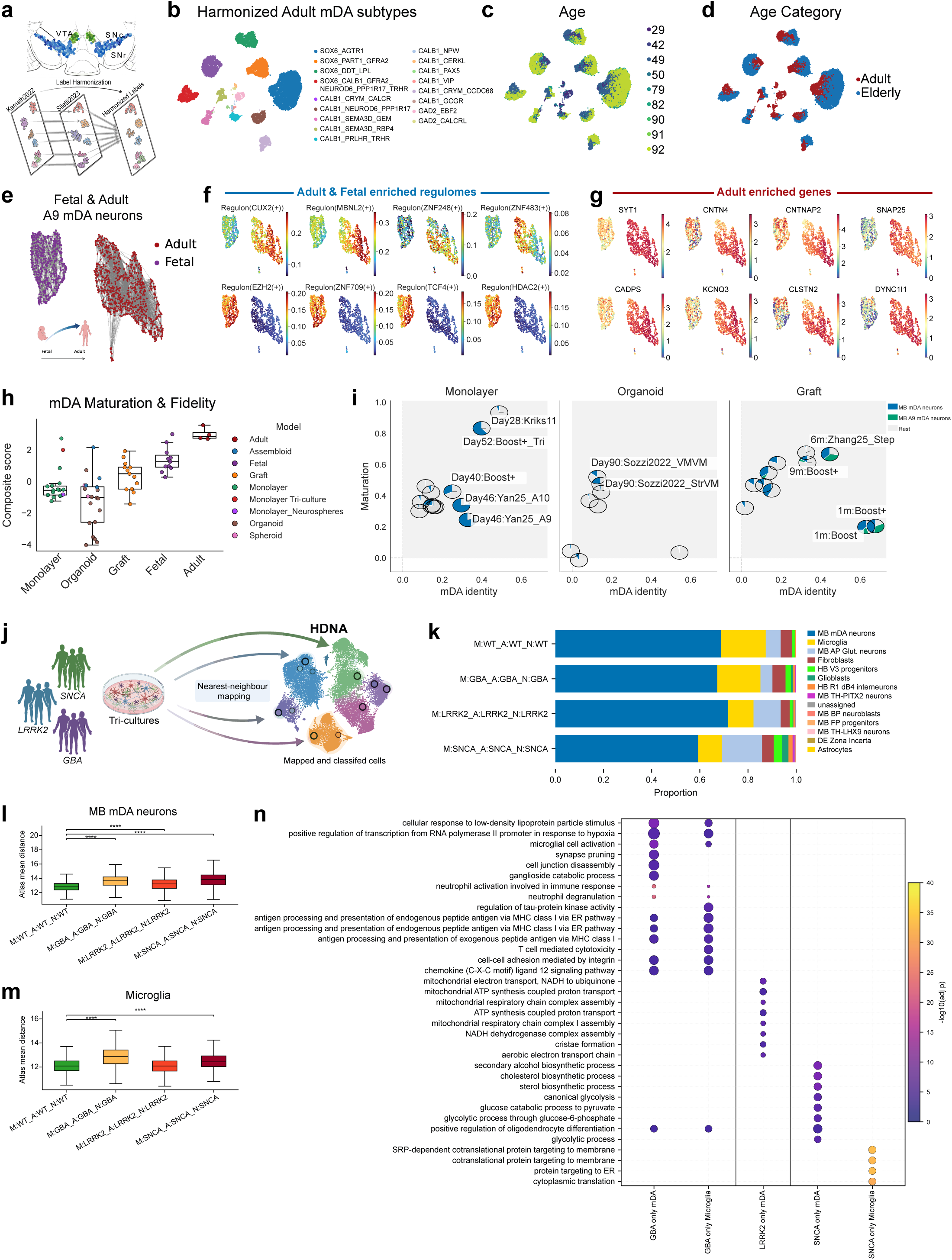
Adult consensus atlas of dopaminergic neurons benchmarks maturation and fidelity across protocols and in Parkinson’s disease tricultures. **a,** Schematic representation of mDA subtype harmonization between post-mortem studies. **b-d**, UMAP coloured by harmonised labels of adult mDAsubtypes (b), by age (c) and by broad age category(d). **e**, UMAP of integrated fetal and adult A9 mDA neurons. **f,** Regulomes identified with SCENIC that enrich in fetal and adult mDA neurons. **g**, UMAP coloured by genes in adult enriched gene modules. **h**, Composite score of dopaminergic identity and maturation across differentiation models. Each point represents a timepoint–protocol–publication group, with the composite score calculated as the sum of z-scored median mDA identity and maturation scores computed across groups. Only protocols with at least 10 mDA neurons were considered. **i**, Dopaminergic identity and maturation across differentiation models with mDA neuronal composition. Each point represents a differentiation condition defined by timepoint, protocol, and study, positioned by median mDA identity (x-axis) and maturation score (y-axis). Points are shown as pie charts indicating the fraction of mDA neurons and A9 mDA neurons, calculated from cells beyond day 25 of differentiation. **j**, Schematic illustrating how additional scRNA-seq datasets from the community can be projected onto the HDNA to extend the and atlas and model PD. **k**, Stacked bar plot showing the proportional composition of cell states across triculture genotypes after projection onto the Human Dopaminergic Neural Atlas (HDNA). **l-m**, Box plots showing the distribution of atlas mean distance for cells annotated as mDA neurons (l) and for microglia (m) after projection onto the Human Dopaminergic Neural Atlas (HDNA). Statistical significance was assessed using one-sided Mann–Whitney U tests comparing each PD genotype to the wild-type control, followed by Benjamini–Hochberg false discovery rate correction. Brackets indicate significant comparisons and stars denote adjusted significance levels (**** q < 0.0001). **n,** Dot plot showing enriched Gene Ontology Biological Process terms for genotype-specific differential-expression signatures from mDA neurons and microglia.

Maturation scoring revealed that most monolayer and organoid systems correlate more with developmental states, with a subset showing increased similarity to adult signatures. Notably, maturation *in vitro* was not strictly time-dependent, as extended culture durations did not consistently result in more adult-like transcriptional profiles (Extended Data Fig. 10e). In contrast, grafted neurons showed the highest correlation with adult signatures, supporting the role of the *in vivo* environment and extended timeframes in promoting maturation, although variability across grafts persisted and were not solely dependant on time (Extended Data Fig. 10e). To quantify the relationship between mDA maturation, fidelity, and cellular stress across the HDNA cohort, we scored single-cell enrichment for glycolysis, hypoxia, and ER stress gene programs alongside mDA identity and adult-associated modules and TFs (Extended Data Fig. 10f). Grafted cells showed the highest mDA fidelity and maturation, coupled with the lowest stress signatures. In contrast, most monolayer and organoid systems exhibited elevated stress and reduced dopaminergic identity, with organoids showing the strongest stress signatures. Notably, tri-culture conditions represented an exception, combining lower glycolytic stress with high mDA fidelity and maturation, suggesting that co-culture with glial components mitigates stress and supports dopaminergic specification even at late timepoints in culture (Fig. 6h, Extended Data Fig. 10f). Overall, stress and metabolic signatures were inversely correlated with mDA identity, but not maturation, indicating that elevated cellular stress is associated with impaired dopaminergic fidelity *in vitro*^7^ (Extended Data Fig. 10g).

We next integrated mDA identity and maturation into a unified ranking framework to identify the best-performing protocols and models, while accounting for conditions that generated sufficient mDA neurons for robust evaluation (Fig. 6h,i). Broadly, graft-derived cells showed the highest overall fidelity and maturation among *in vitro* systems, approaching fetal mDA neurons, whereas monolayer and organoid models scored lower overall, with organoids displaying the greatest variability across conditions (Fig. 6h). In monolayer cultures, the highest scores combined with a high percentage of mDA neurons were achieved by Boost+^23^ under triculture conditions and the Yan25 A9/A10^21^ protocols (Fig. 6i). In 3D systems, the highest-scoring conditions were predominantly assembloid models, suggesting that co-culture with target cell types enhances dopaminergic fidelity and maturation (Fig. 6i). Among grafted mDA neurons, the Zhang25 stepwise^22^ and Boost+^23^ protocols emerged as the top-ranked conditions (Fig. 6i). Collectively, these results establish dopaminergic identity and maturation as distinct, partially independent dimensions and demonstrate how this framework enables systematic benchmarking of differentiation conditions.

Given that triculture systems combine high dopaminergic identity and maturation with robust mDA neuron yield, they represent an optimal platform for disease modeling. We therefore leveraged this system to test the utility of the HDNA framework in interrogating PD–associated perturbations. We generated iPSC-derived tricultures^86^ carrying PD mutations (GBA, LRRK2 or SNCA)^2,71–74^, in which all cell subtypes shared the same genotype (see methods) and then developed a toolkit for projecting new datasets onto the HDNA (Fig. 6j). Single-cell profiles were mapped onto the HDNA to assign identities and quantify subtype composition. Most cells were mDA neurons and microglia, with only very few astrocytes (Fig. 6k), likely reflecting the initial mixing ratio (8 mDA neurons:2 microglia:1 astrocytes), suggesting that increasing astrocyte input may improve representation. PD-mutant mDA neurons showed increased divergence from the HDNA reference (Fig. 6l), while GBA and SNCA microglia, but not LRRK2, also deviated from controls (Fig. 6m), indicating mutation-specific effects across lineages. Finally, differential expression analysis combined with pathway analysis revealed distinct genotype- and cell-specific responses (Fig. 6n, Supplementary Table 7). GBA mDA neurons and microglia were enriched for immune-interaction related pathways, consistent with the growing literature linking GBA-associated PD to neuroinflammatory phenotypes^87^. Instead, LRRK2 mDA neurons enriched to mitochondrial electron transport pathways, in line with the established role of LRRK2 in mitochondrial dysfunction^88,89^. In contrast, SNCA mDA neurons showed enrichment for cholesterol and sterol biosynthesis, supporting a link between α-synuclein pathology and lipid metabolic dysregulation^90–93^. SNCA microglia, meanwhile, were enriched for cotranslational protein targeting to membranes, suggestive of altered proteostasis^94^. Together, these analyses demonstrate that the HDNA provides a unified framework for benchmarking stem-cell derived PD models and identifying genotype- and cell-type–specific molecular perturbations.

## DISCUSSION

In this study, we establish an integrated framework to benchmark human stem cell–derived mDA differentiation protocols against high-resolution developmental and adult human brain atlases, enabling quantitative assessment of dopaminergic fidelity, maturation, and protocol-specific off-targets. Using CapybaraBrain, we show that, relative to fetal cells, *in vitro* cells are less dominated by a single cell-state program and more frequently exhibit hybrid identities, indicating incomplete lineage resolution and weakened transcriptional boundaries compared with fetal development. These findings support a model in which the Waddington landscape^95,96^ is shallower and less canalized *in vitro* than *in vivo*, causing weaker fate commitment and increased occupancy of hybrid cell states.

We observe substantial heterogeneity in dopaminergic yield and identity across protocols and models. Grafted cells show the highest fidelity to midbrain dopaminergic programs and exhibit A9-like features, indicating that *in vivo* environments promote subtype specification. In contrast, all *in vitro* systems, generate dopaminergic neurons with lower fidelity and fail to robustly produce A9-like neurons, suggesting they lack key cues for this fate in culture. Beyond differences in on-target yield, our analysis reveals recurrent off-target subtypes that reflect systematic patterning errors. A prominent example is the TH-PITX2 neuronal subtype, a related but distinct identity from canonical mDA neurons, placed by mouse anatomical analysis in the rostral midbrain and posterior hypothalamus.

A second recurrent off-target population are hindbrain V3 progenitors, detected across multiple protocols where they often persist even after grafting, marking them as a clinically relevant contaminant. These progenitors rarely progress toward serotonergic derivatives, remaining locked in a ventral floor plate-like state with limited neurogenic competence^75^, likely due to lack of *NKX2-2* expression^76,77^. These findings may explain the relative paucity of serotonergic neurons in most protocols and models. Instead, these progenitors are biased toward gliogenic fates, as validated by genetic fate mapping in mouse models showing that hindbrain, but not midbrain, floor plate cells carry this competence. Together, these findings indicate that most protocols strongly induce ventral floor plate identity via SHH signalling, but fail to maintain precise rostrocaudal patterning, allowing drift toward either hypothalamic or hindbrain floor plate domains.

This positional instability co-occurs with a further recurrent feature: fibroblast-like transcriptional programs within ventral progenitor populations, marked by extracellular matrix genes and EMT-associated factors, and are enriched in monolayer cultures. Our *in vivo* fate-mapping experiments show that ventral Gli1 progenitors of both midbrain and hindbrain can generate perivascular fibroblast populations. This suggests that non-neuronal cells in stem cell-derived grafts arise from culture-induced mechanical and cell-matrix cues^97–99^ acting across multiple ventral progenitor populations, rather than midbrain floor plate alone^17^. Critically, these relationships were only resolvable through CapybaraBrain, which detects latent and nested lineage potentials and early off-target trajectories that single-label classifiers would miss.

A further contribution of this study is the integration of developmental and adult references to disentangle dopaminergic identity from maturation. By harmonizing adult post-mortem mDA subtype datasets^52,53^ into a consensus atlas, we provide a common framework for assessing how closely *in vitro*-derived cells approach mature states and show that mDA identity and maturation were not fully coupled. This distinction is important for interpreting both graft quality and disease-model relevance. Our data further show that elevated glycolysis, hypoxia and ER stress signatures are strongly associated with lower mDA fidelity but not maturation, especially in 3D models^7,23^. Notably, tricultures exhibit reduced stress alongside higher fidelity and maturation, suggesting glial incorporation as an actionable strategy for improving dopaminergic differentiation. Finally, we provide a framework to continuously expand the HDNA and apply it to disease modelling. By projecting new PD tricultures onto the atlas, we illustrate how disease-associated mutations can be evaluated against a common reference, allowing both compositional benchmarking and cell-type-specific measurement of transcriptional divergence. This will facilitate future studies aimed at separating disease effects from protocol-specific variation and comparing PD models generated across different laboratories and platforms.

We anticipate that these atlases will serve as a gold-standard resource for the field, enabling standardized comparison of existing and emerging protocols, improving interpretation of graft composition, and accelerating refinement of stem cell–derived models for PD and cell therapy. More broadly, the framework established here can extend beyond dopaminergic neurons by enabling rigorous assessment of stem cell products, exposing hidden off-target states early in differentiation and revealing systematic compositional biases across cell populations. Since our developmental atlas spans diverse neuronal lineages, this strategy can be readily applied to other stem cell–derived neural models, including serotonergic, noradrenergic and cerebellar systems to model specific diseases^100–107^. In parallel, the integrated adult post-mortem atlas provides a benchmark not only for mDA subtype fidelity, but also for emerging approaches aimed at promoting maturation^83,108,109^ and aging^110–115^ in hPSC models. Collectively, our study provides a generalizable roadmap for evaluating cell-subtype fidelity, detecting aberrant trajectories, and rationally engineering more faithful human brain models across regions and diseases.

## METHODS

### Human fetal sample acquisition

For our unpublished developmental droplet and spatial transcriptomics data, human developmental brain tissue samples <12 PCW were obtained from elective terminations under REC 96/085 (East of England - Cambridge Central Research Ethics Committee). Samples > 12 PCW were obtained from the MRC/Wellcome Trust Human Developmental Biology Resource under full ethical approval from the London-Fulham Research Ethics Committee. Informed and written consent was obtained for the acquisition of all samples. Briefly, samples were kept suspended in PBS and at - 4°C on ice during dissection. Brainstem regions were separated according to gross morphology under direct visualization using a microscope. Approximations were made to capture the entirety of the midbrain region (< 12PCW). For older samples (>12 PCW), midbrain dissections were performed by the HDBR team. Following dissection, samples were embedded in optimal cutting temperature medium (OCT) and frozen at -80°C on an isopentane-dry ice slurry.

### Human fetal sample processing for single-nuclei RNA and ATAC sequencing

For nuclei dissociation, cryosections of the frozen blocks were cut at a thickness of 10 μm using a Leica CM1950 cryostat. A tissue homogeniser was then applied in homogenization buffer (250 mM sucrose, 25 mM KCl, 5 mM MgCl2, 10 mM Tris-HCl, 1 mM dithiothreitol, 1× protease inhibitor, 0.4 U μl−1 RNaseIn, 0.2 U μl−1 SUPERaseIn and 0.1% Triton X-100 in nuclease-free water), as described previously^116^, to dissociate and filter for nuclei. 7 AAD-staining was performed, followed by cell sorting using a BigFoot spectral cell sorter (ThermoFisher Scientific) and proprietary software. Gating for 7AAD was performed to identify intact nuclei and remove cell debris. Viable nuclei were processed in accordance with the 10X Genomics Multiome ATAC + RNA protocol. Droplets were loaded for a targeted recovery of 16,000 per reaction where possible. Quality control of cDNA libraries were done using an Agilent bioanalyzer. Libraries were sequenced on a NovaSeq 6000 platform (Illumina) with a minimum depth of 20,000 paired reads per droplet. For this study only the RNA portion of the data was used.

### Mouse ventral midbrain tissue *in situ* hybridization

Four C57BL/6 mice (JAX#000664; 1.5-2.1 months old; two females and two males) were deeply anesthetized and transcardially perfused with phosphate-buffered saline (PBS). Brains were rapidly harvested, flash-frozen in FSC 22 Frozen Section Media (Leica Biosystems, #3801480), and stored at -80C. Coronal sections were cut at 16 μm thickness on a cryostat (Leica Biosystems, CM3050S), mounted onto SuperFrost Plus Microscope Slides (Fisherbrand, #1255015), and kept at -80C until use. RNAscope assay was performed on fresh frozen brains, using the RNAscope Multiplex Fluorescent Reagent Kit v2 (Advanced Cell Diagnostics). The probes used were Mm-Slc17a6, Mm-Pitx2-C2, and Mm-Th-C3 RNAscope probes (Advanced Cell Diagnostics; #319171, 412841-C2, and 317621-C3). Prior to hybridization, sections were post-fixed with 4% paraformaldehyde (PFA) in PBS for 2 hours at room temperature (RT). Slides were then dehydrated via an ethanol series (50%, 70%, 100%, 100%) and air-dried. Sections were then treated with hydrogen peroxide for 10 min at RT, rinsed with distilled water, treated with Protease Plus for 10 min at RT, and rinsed in PBS. Sections were incubated with the probe mixture for 2 hours at 40°C in a HybEZ oven (Advanced Cell Diagnostics). Probes were diluted at a 50:1:1 ratio (C1:C2:C3). After hybridization, signal amplification was performed (AMP1 for 30 min; AMP2 for 30 min; AMP3 for 15 min). Next, each channel’s signal was developed by treating sections with appropriate HRP reagents for 15 min at 40°C, followed by Opal fluorophore reagents for 30 min at 40°C (Opal 690 (FP1497001KT) for C1, Opal 570 (FP1488001KT) for C2, Opal 520 (FP1487001KT) for C3, 1:750 dilution in TSA buffer; Akoya Biosciences). HRP activity was blocked using HRP blocker for 15 minutes at 40°C. Sections were washed with Wash Buffer between all incubation steps. Finally, sections were stained with DAPI and mounted with ProLong Gold Antifade Mountant (Invitrogen, Cat#P36930). Step-by-step methods can be found at protocols.io (10.17504/protocols.io.36wgqp83kvk5/v1). Fluorescent images were acquired using an Olympus BX61VS slide scanner (10x objective lens) and a ZEISS LSM 900 confocal microscope (20x and 63x objective lens) for high-resolution analysis. Confocal images were acquired as Z-stacks (10 slices, 0.49 μm step size for 20x; 15 slices, 0.21 μm step size for 63x). All images within a single set of experiments were acquired with identical acquisition settings, including laser power, exposure time, detector gain, digital offset, and scan speed, for each respective wavelength. Figures were prepared using Adobe Illustrator. Slc17a6, Pitx2, and Th positivity was determined quantitatively by counting the number of puncta surrounding a cell’s nucleus. A cell was classified as positive if it exhibited six or more distinct, clustered puncta corresponding to the C1, C2, or C3 signal.

### Genetic inducible fate mapping

Mouse lines: All experiments conducted on mice were performed in accordance with the welfare animal regulations of the Federal German Government, European Union and were approved by the LAVE NRW. All animals were housed in a controlled environment with 12-h light/night cycles and had free access to food and water. The following mouse lines were used: *Shh^CreER^* mice (Shhtm2(cre/ERT2)Cjt/J, JAX 005623), *Gli1^CreER^* mice (Gli1tm3(cre/ERT2)Alj/J, JAX 007913), *R26^Ai9^* mice (Gt(ROSA)26Sortm9(CAG-tdTomato, JAX 007909) and *Aldh1l1-GFP* mice (Tg(Aldh1l1-EGFP)OFC789Gsat/Mmucd, RRID:MMRRC_011015-UCD, obtained from MMRRC at University of California at Davis, a strain repository funded by NIH. This strain was donated to the MMRRC by Nathaniel Heintz, Ph.D., The Rockefeller University, GENSAT). All mice were maintained in an outbred CD1 background. Both male and female mice were used for fate mapping experiments. To facilitate the detection of astrocytes in Shh- or Gli1-genetic inducible fate mapping (GIFM) experiments, *Shh^CreER/+^, R26^Ai9/Ai9^* or *Gli1^CreER/+^, R26^Ai9/Ai9^* mice were crossed with *Aldh1l1-GFP* mice. Noon of the day that a vaginal plug was detected was designated as E0.5. Tamoxifen (TM; T-5648 Sigma, St. Louis MO, USA) was dissolved in corn oil (Sigma C8264) and administered by oral gavage (100 mg/kg body weight) to pregnant females.

#### Tissue processing

Adult *Shh^CreER/+^, R26^Ai9/+^, Aldh1l1-GFP* or *Gli1^CreER/+^, R26^Ai9/+^, Aldh1l1-GFP* mice were perfused transcardially with phosphate buffered saline (PBS) followed by 4% paraformaldehyde (PFA). Dissected brains were post-fixed in PFA overnight at 4°C, cryoprotected in sucrose, embedded in Tissue Tek and sectioned into 30 µm thick free-floating cryosections.

#### Immunofluorescence staining

Sections were rinsed in PBS and PBS + 0.1% TritonX-100 (0.1 PBT) and then blocked in 0.1PBT + 10% NDS for 2 h at RT. Sections were incubated with primary antibodies in 0.1 PBT + 3% NDS overnight at 4°C. The following primary antibodies were used: rabbit anti-COL1A1 (1:200, Abcam, ab34710), rat anti-GFP (1:500, Nacalai Tesque, 04404-84), chick anti-RFP (1:500, Novus Biologicals, NBP1-97371) and sheep anti-TH (1:500, Sigma-Aldrich, AB1542). For the COL1A1 antibody, the blocking step was preceded by antigen retrieval (Tris-EDTA pH8 for 30 min at 80 °C). After washing in 0.1 PBT, sections were incubated with secondary antibodies in 0.1 PBT + 3% NDS for 2 h at RT. The following secondary antibodies (all donkey-derived; from Thermo Fischer Scientific) were used at a dilution of 1:1000: anti-chicken-Alexa 555 (A78949), anti-rabbit Alexa 647 (A31573), anti-rat-Alexa 488 (A21208), anti-sheep Alexa 647 (A21448). Hoechst 33258 (Abcam) was used to counterstain nuclei.

#### Imaging

Images were acquired with a Spinning-disk confocal microscope (Visitron Visiscope, Visitron Systems) using standard excitation and emission settings for 405, 488, 561 and 647 nm signals with 10X (Plan-Apochromat, 10x/0.45) and 63X (water immersion, C-Apochromat, 63x/1.2) lenses. Tile images were stitched using VisiView software (Version 6.0.0.37, Visitron Systems).

#### Detection of fate-mapped astrocyte and fibroblast

Single-plane images of GFP, RFP and TH immunostaining were captured with a 10X lens. Shh-GIFM: Every eighth coronal section (corresponding to 240 µm intervals) was evaluated in the mid/hindbrain region. For quantification, posterior midbrain sections at the level of the retrorubral field and anterior hindbrain sections at the level of the locus coeruleus were evaluated (1-2 sections per region, n=3 mice). To detect fate-mapped astrocytes, Cellpose (Version 3.1.0) was used to segment GFP+ cell somata to generate regions of interest (ROI). RFP+ ROIs were identified with Fiji (ImageJ version 1.54f) using a custom script based on image specific background thresholds. ROIs were manually curated as needed. To quantify fate-mapped astrocytes, the selected ROIs were converted to masks (Fiji “Masks from ROIs”, Version 1.0.0^117^) and analysed using “Analyze Particles” (Fiji). Gli1-GIFM: to detect the distribution of fate mapped fibroblasts (RFP+/COL1A1+ cells), Cellpose was used to segment nuclei of COL1A1+ cells and to generate ROIs. RFP+ ROIs were selected as described for fate-mapped astrocytes. To visualize the distribution of co-expressing cells, fate-mapped fibroblasts were marked in red and fate-mapped astrocytes in orange using Fiji. N=3 mice.

### Tri-culture system, co-culture ratios and single-cell analyses

Experiments involving human pluripotent stem cells (hPSCs) were approved by the Tri-institutional ESCRO committee. hPSCs were maintained on Vitronectin-coated plates in Essential 8 medium as previously described^118^ . GBA-N370S and LRRK2-G2019S and SNCA A53T mutations were introduced into the H9 parental line using prime editing^119,120^. Briefly, pegRNAs containing the respective spacer and 3′ extension sequences were cloned into the pU6-tmpknot-GG-acceptor backbone (Addgene #174039). The corresponding nicking sgRNAs were cloned into a nicking sgRNA backbone (Addgene #47108). To create the mutation induction, the PEmax-P2A-hP53DD (Addgene #214084), epegRNA, and nicking sgRNA plasmids were delivered into H9 cells via electroporation, followed by single-cell cloning. Target loci from individual clones were amplified by PCR and analyzed by Sanger sequencing to identify correctly edited clones. The sequences used for cloning were as follows: For GBA-N370S induction, the pegRNA spacer sequence was AGCCGACCACATGGTACAGG, the 3′ extension was TTACCCTAGAGCCTCCTGTACCATGTGGTC, and the nicking sgRNA spacer sequence was ACCCTTACCTACACTCTCTG. For LRRK2-G2019S induction, the pegRNA spacer sequence was ATTGCAAAGATTGCTGACTA, the 3′ extension was AGCAATGCTGTAGTCAGCAATCTTTGC, and the nicking sgRNA spacer sequence was GACAGACCTGATCACCTACC. For SNCA A53T induction, the pegRNA spacer sequence was GGAGGGAGTGGTGCATGGTG, the 3′ extension was ACCTGTTGTCACACCATGCACCACT, and the nicking sgRNA spacer sequence was TCATAGGAATCTTGAATACT. Dopamine neuron–astrocyte–microglia tri-cultures were generated from isogenic WT or mutant hPSC lines carrying GBA N370S, LRRK2 G2019S or SNCA A53T. Dopamine neurons^23^ were purified at day 25 by CD49e magnetic depletion and plated into poly-L-ornithine/fibronectin/laminin-coated 96-well plates at 112,000 cells per well. Microglia^86^ and astrocytes^118^ were dissociated and added at 28,000 and 14,000 cells per well, respectively, yielding a neuron:microglia:astrocyte ratio of 8:2:1. Cultures were maintained in NB/N2/B27 medium supplemented with BDNF, GDNF, ascorbic acid, cAMP, IL-34, and M-CSF as previously described^86^. At day 9, cells were dissociated using the Worthington Papain Dissociation System and labeled with TotalSeq hashtag antibodies for single-cell RNA sequencing. Libraries were prepared using the Chromium Single Cell 3′ v3.1 platform (10x Genomics), targeting ∼30,000 cells per run. Reverse transcription, barcoding, cDNA amplification, and library preparation followed the manufacturer’s protocols. Samples were multiplexed using Hash Tag Oligonucleotides as previously described^121^ and sequenced on an Illumina NovaSeq. Mapping was performed using the human reference genome GRCh38 with CellRanger. HTO sequencing data were aligned to the HTO barcodes, and UMIs were counted for each cell using CITE-seq-Count. Using a two-component K-means algorithm, we partitioned logged HTO counts into two distributions: background noise (lower mean) and positive tags (larger mean). Each droplet was then assigned to its source sample based on tags in the positive signal component. We classified droplets with multiple assignments as doublets and those with a single assignment as singlets. This analysis was performed using SHARP v0.1.1.

### hPSC culture and midbrain dopaminergic differentiation for datasets from the Kunath Lab

Human ESC lines MasterShef7 (MShef7) and RC17 and human iPSC lines 1231A3 and 404C2 were maintained under feeder-free conditions. hESCs were cultured on Laminin-521 in StemMACS iPS-Brew XF medium, and iPSCs on iMatrix-511 silk in StemFiT AK-02N medium. All differentiation strategies were based on ventral midbrain floor plate induction using dual SMAD inhibition combined with SHH and WNT pathway activation. Three related protocols were used.

#### Kriks11^122^ based

hESCs and iPSCs were seeded at 40,000 cells/cm^2^ onto Laminin-111 -coated plates, and differentiated using dual SMAD inhibition (SB431542 and LDN193189) with SHH-C24II and CHIR99021 in a 1:1 DMEM/F12:Neurobasal medium supplemented with B27 (minus vitamin A), N2 and L-glutamine. Y-27632 was included during the first 48h only. CHIR99021 concentration (0.9-1.1 μM) was titrated per cell line. From day 4 onward, patterning conditions were maintained with 50% reduced N2/B27, and FGF8b and heparin were introduced at day 9 to reinforce midbrain identity. At day 11, cells were dissociated with Accutase and replated at 800,000 cells/cm2 in Neurobasal-based medium containing FGF8b, heparin, BDNF, GDNF, and ascorbic acid. A second replating was performed at day 16, after which cells were maintained in terminal differentiation medium containing BDNF, GDNF, ascorbic acid, dbcAMP and DAPT. A detailed version of this protocol is available at: dx.doi.org/10.17504/protocols.io.bddpi25n.

#### Kyoto^6^ protocol (JP)

The cells were differentiated in the Morizane Lab, and then the frozen cell vials were sent to the Kunath Lab. Human iPSCs were seeded at 500,000 cells/cm^2^ onto iMatrix-511 silk-coated plates and differentiated in medium supplemented with KnockOut Serum Replacement (Thermo Fisher Scientific). Ventral midbrain identity was specified using dual SMAD inhibition (LDN193189 and A83-01) in combination with SHH pathway activation (purmorphamine), FGF8, and canonical WNT activation (CHIR99021), with the medium refreshed daily. At day 11, cells were dissociated with Accumax and replated at 500,000 cells/cm² onto iMatrix-511 silk–coated plates, and cultures were maintained in neural differentiation medium with ascorbic acid, BDNF, GDNF, with dbcAMP and DAPT added from day 21 to promote maturation. A detailed original protocol is available at: doi: 10.1016/j.stemcr.2014.01.013.

#### Boost^24^ based protocol

The CHIR protocol was based on the Kriks11 protocol with a transient increase in WNT signaling. CHIR99021 concentration was increased to 2.5 μM for MShef7 and to 7.5 μM for RC17 from day 4 to day 7 (Kim et al, 2021).

#### Single-cell RNA sequencing and sample multiplexing

Single-cell transcriptomic profiling was performed using the Chromium Next GEM Single Cell 3′ v3.1 platform (10x Genomics) in combination with the 3′ CellPlex sample multiplexing system. Individual samples were labelled with lipid-conjugated Cell Multiplexing Oligos (CMOs; 305-312), each containing a unique 15-nucleotide Feature Barcode sequence to permit post hoc demultiplexing. For each condition, 1 x 10^6^ cells were incubated with the assigned CMO in a 100 µL staining reaction, followed by two washes in PBS containing BSA. DAPI-negative cells were purified by flow cytometry to enrich for viable cells. Following labelling, samples were pooled in equal proportions, with approximately 15,000 cells contributed per condition. Four uniquely barcoded samples (total ∼60,000 cells) were combined per Chromium channel, allowing multiplexed analysis of Kriks11, JP, Boost and self-renewing (SR) conditions within each sequencing run to minimise batch effects. Across eight Chromium libraries, this strategy permitted analysis of 32 distinct experimental conditions. Approximately 60,000 cells were loaded per channel to achieve high recovery while accounting for Poisson-based encapsulation. Gel Bead-in-Emulsion (GEM) generation resulted in simultaneous capture of polyadenylated mRNA and CMO-derived Feature Barcode molecules within individual droplets. Following reverse transcription, cDNA was amplified (11-13 PCR cycles depending on run), and size selection was performed to separate gene expression and multiplexing products. For each channel, two libraries were generated: a 3′ gene expression (GEX) library with an average fragment size of approximately 450 bp, and a CellPlex Feature Barcode library with an average fragment size of approximately 200 bp. Libraries were dual indexed and quantified using High Sensitivity DNA assays prior to sequencing. Eight GEX libraries and eight corresponding multiplexing libraries were submitted for sequencing. Sequencing was performed on an Illumina NextSeq 2000 using a P3 flow cell and 100-cycle configuration, following 10x Genomics recommendations for Chromium Single Cell 3′ v3.1 chemistry. Sequencing depth targets were at least 20,000 read pairs per cell for gene expression libraries and 5,000 read pairs per cell for multiplexing libraries. Raw data were processed with Cell Ranger *multi* to demultiplex pooled libraries into the original samples. CMO barcode features were then removed from the gene list prior to downstream analysis. Initial quality control was performed in R by excluding outlier cells identified with *scater::isOutlier()*. Raw sequencing data have been deposited in the European Nucleotide Archive (ENA) under accession number ERP191407.

### hPSC culture and midbrain dopaminergic differentiation for datasets from the Croft Lab

Human pluripotent stem cells (H9) were cultured on lamin521(biolamina) in StemFlex(Thermo) as previously described^123,124^ Midbrain differentiation was performed per^23,125^ with the following changes: PSC spheres were aggregated with uniform size following^126^, and cultured to day 0-33 with no dissociation, no DAPT was used, 1% Albumax was added throughout differentiation, CHIR99021 was administered in a modified biphasic regimen adapted from^23^: 1 µM (days 0–3), 6 µM (days 4–6), and 3 µM (days 7–11) and SHH protein was replaced with 1uM SAG.13 and 1uM Purmorphamine, as described^127^. Organoids were dissociated at day 33 with Papain (Worthington, following adapted protocol^128^), and cryopreserved. For single cell captures, FPP were thawed, depleted for dying cells with AnnexinV microbeads (Miltenyi, Dead cell removal kit), and debris was floated with a density cushion (4% BSA in L15, adjusted to pH7 CITE). Cells were hashtagged, pooled, and captured on 10x Chromium 3’ Gene Expression. The libraries were subsequently sequenced to an average of ∼20K paired-end reads/cell following standard methods. Raw read quality control, alignment and quantification was performed by Cell Ranger (v7.1.0). Cell barcodes were demultiplexed into their cell line of origin using a combination of antibody hashtagging (Biolegend) and isogenomic SNP data reference using *Demuxafy*^129^ (v3.0.0). Quality control was performed to remove poor-quality cells, doublets, and unassigned cells.

### hPSC culture and midbrain dopaminergic differentiation for datasets from the Parmar Lab

#### Human pluripotent stem cell culture

Undifferentiated hPSC lines RC17 (sourced by Roslin Cells, cat. no. hPSCreg RCe021-A), and TH-Cre knock-in line (clone B2)^130^ were maintained on 6-well plates (Sarstedt, cat. no. 83.3920) coated with recombinant laminin-521 (Biolamina, 0.5 μg/cm2 in DPBS+Mg^2+/^Ca^2+^) and cultured in iPS Brew XF medium (Miltenyi Biotec, GMP-grade). At 70-90% confluence, cells were passaged using EDTA (0.5 mM for 7 min at 37 °C) and reseeded at 5k-10k cells/cm² in iPS Brew supplemented with 10 μM ROCK inhibitor (Y-27632, Miltenyi, cat. no. 130-106-538) for the first 24 hours.

#### Ventral midbrain organoid differentiation

Ventral midbrain (vMB) organoids were generated as described in^19^. Prior to differentiation, hPSC cultures at 70–90% confluency were dissociated into single-cell suspensions using Accutase (50 μL/cm², Thermo Fisher Scientific, cat. no. A1110501) for 5 min at 37°C. Aggregates were obtained by seeding 8k cells/well in U-bottom 96-well plates (Corning, cat. no. 7007) in iPS-Brew supplemented with 10 μM Y-27632. Differentiation medium (1:1 DMEM/F12 and Neurobasal) was supplemented with N2 (1:100), SB431542 (10 μM, Miltenyi, cat. no. 130-106-543), rhNoggin (100 ng/mL, Miltenyi, cat. no. 130-103-456), SHH-C24II (400 ng/mL, Miltenyi, cat. no. 130-095-727), and CHIR99021 (1.5 μM for RC17 and TH-Cre lines, Miltenyi, cat. no. 130-106-539), alongside L-glutamine (2 mM), MEM-NEAA (1×), penicillin-streptomycin (20 U/mL), and 2-mercaptoethanol (50 μM). FGF-8b (100 ng/mL, Miltenyi, cat. no. 130-095-740) was introduced from day 8. On day 11, the medium was replaced with Neurobasal with B27 (–vitamin A, 1:50), BDNF (20 ng/mL, Miltenyi, cat. no. 130-096-286), L-ascorbic acid (200 μM, Sigma-Aldrich, cat. no. A4403), and FGF-8b. At day 16, vMB organoids were embedded in 30 μL Matrigel droplets (Corning, cat. no. 354234) together with another vMB or STR organoids to form assembloids. Terminal differentiation medium consisted of Neurobasal with B27 (–vitamin A), BDNF (20 ng/mL), L-ascorbic acid (200 μM), db-cAMP (500 μM, Sigma-Aldrich, cat. no. D0627), GDNF (10 ng/mL, R&D Systems, cat. no. 212-GD-010), and DAPT (1 μM, Tocris Bioscience, cat. no. 2634), maintained for up to four months.

#### Striatal organoid differentiation

STR organoids were generated following the protocol described in^131^ with minor modifications. hPSCs (RC17) were seeded in U-bottom 96-well plates as described above and cultured for the first 6 days in differentiation medium (1:1 DMEM/F12 and Neurobasal, supplemented with N2 1:100) containing SB431542 (10 μM, Miltenyi, cat. no. 130-106-543) and rhNoggin (200 ng/mL, Miltenyi, cat. no. 130-103-456). From day 6 to day 16, organoids were maintained in Neurobasal medium supplemented with IWP-2 (2.5 μM, Selleck Chem, cat. no. S7085) and Activin A (100 ng/mL, Peprotech, cat. no. 120-14E) to induce rostral fate. From day 12, the retinoid X receptor agonist SR11237 (100 nM, Tocris, cat. no. 3411) was also included. From day 16, STR organoids were fused with vMB organoid as described above and maintained in the same terminal differentiation medium as vMB organoids.

#### Sample preparation for single cell RNA sequencing

Assembloids at day 120 of differentiation were washed three times with DPBS (CTS, Thermo Fisher Scientific, cat. no. A1285601) and dissociated to a single cell suspension using a papain and DNase I solution in EBSS (Earle’s Balanced Salt Solution) from the Papain Dissociation kit (Worthington, cat. no. LK003150). A total of 7k - 10k cells per sample were used as input for scRNAseq library preparation.

#### scRNA-seq library preparation and sequencing

Single-cell suspensions were loaded onto Chromium Single Cell 3′ Chips (10X Genomics, cat. no. PN-120233, v3.1 chemistry) along with the master mix following the manufacturer’s instructions to generate single-cell gel beads in emulsion (GEMs). Library quality was assessed by Bioanalyzer (DNA HS kit, Agilent) prior to sequencing. Libraries were sequenced on an Illumina NovaSeq 6000 in 2×100 bp paired-end mode. Raw base calls were demultiplexed using bcl2fastq (v2.19) and processed through the Cell Ranger pipeline (10X Genomics, v3.0) with alignment to the GRCh38 reference genome to generate count matrices for downstream bioinformatic analysis.

### Boost+ Immunohistochemistry for TH-PITX2 validation

Human pluripotent stem cells (KOLF2.1) were differentiated using the Boost+ protocol^23^ . Cells were fixed in 4% paraformaldehyde (PFA; Affymetrix) prepared in DPBS for 13 minutes at room temperature, followed by washes in DPBS. Samples were then permeabilized with 0.5% Triton X-100 in DPBS for 25 minutes and blocked for 1 hour in blocking buffer containing 2% BSA and 0.25% Triton X-100 in DPBS. Primary antibodies against PITX2 (R&D, AF7388), TH (Sigma-Aldrich, MAB318), and FOXA2 (Cell Signaling, 8186S) were diluted in blocking buffer and applied overnight at 4 °C. After washing with DPBS, samples were incubated for 1 hour at room temperature with Alexa Fluor-conjugated secondary antibodies (488, 555, or 647; Invitrogen) diluted in blocking buffer. Following additional DPBS washes, nuclei were counterstained with DAPI (Sigma) for 5 minutes. Images were captured using Olympus or Zeiss inverted fluorescence microscopes.

### Published midbrain fetal integration and label harmonization

All processing and analyses were carried out using scanpy^132^ unless indicated otherwise. Highly variable genes were identified per study using scanpy.pp.highly_variable_genes with the study specified as batch_key. Single-cell datasets from the LaManno^43^ study and the Braun^44^ studies were integrated and their annotations harmonized using CellHint^56^. Cell type alignment across datasets was performed using cellhint.harmonize, specifying the study as the batch variable and the original cell type annotations as input labels. Manual annotation guided by canonical marker selection refined the alignment to create a harmonized label for each cell subtype. The LaManno study was used only for harmonization of labels but was not integrated in the final fetal atlas due to very low number of cells.

### Published diencephalon and hindbrain fetal annotation

Cells corresponding to diencephalic and hindbrain regions were downloaded from the Braun dataset via CellxGene. For the diencephalon, the CellxGene precomputed embedding (X_embedding) was used to guide annotation, and cell identities were assigned based on Leiden clustering (Leiden_1.0) and literature-supported canonical marker genes.

For hindbrain regions, a precomputed embedding was not available in CellxGene. Therefore, cells in dissection regions labelled as Hindbrain, Pons, Cerebellum, or Medulla in the Braun dataset were retrieved and processed de novo. Analysis was restricted to neural lineages by selecting cells classified as Neuron, Radial glia, Neuroblast, or Neuronal intermediate progenitor cells (IPC). A neighbourhood graph was constructed using 30 principal components and 30 nearest neighbours with cosine distance. Batch effects across donors were corrected using BBKNN with DonorID as the batch variable. UMAP embeddings were then computed for visualization. Cell populations were identified using Leiden clustering (Leiden_1.4) and annotated based on literature-supported canonical marker genes. To obtain a unified representation of the integrated dataset, we applied the CellHint integration function cellhint.integrate, using DonorID as the batch variable and the detailed cell type labels as input annotations (n_meta_neighbors = 1) for the diencephalic and the hindbrain datasets independently. The resulting integrated embedding was used for all downstream visualizations.

### New human fetal sample processing and cell classification

Mapping was performed using the human reference genome GRCh38-2020-A on CellRangerArc (v2.0.0) software. Subsequently, the resultant matrices were processed using CellBender^133^ (v2) for ambient RNA profile calculation and removal. The filtered matrix was then analysed using Scrublet^134^ to remove potential doubles (threshold of > 0.3 doublet probability). Briefly, per-droplet scrublet scores were first determined for CellRanger-arc count-matrices from each 10x Multiome (gene-expression) lane independently. The droplets were then overclustered through the standard scanpy workflow using default parameters up to Leiden clustering. Each individual cluster was further clustered. A per-cluster median of scrublet scores were computed. A normal distribution of doublet score, centred at the score median with a standard deviation estimated from the median absolute deviation (MAD) was used to compute p-values for each of the clusters. After false-discovery rate adjustment using Benjamini-Hochberg correction, a p-value of >0.65 was applied as an additional cutoff value. To assign cell identities we trained CellTypist^67^, a logistic regression classifier, on the concatenated and harmonized new labels of the midbrain, diencephalon, and hindbrain of the Braun dataset. To limit class imbalance, the dataset was downsampled across *Detailed_CellTypes* labels, retaining a maximum of 2,500 cells per cell type. Training was performed without feature selection, using a maximum of 500 iterations. Cell identities in the new fetal data were predicted using CellTypist with majority_voting = True, mode = ’prob match’, and a probability threshold of 0.8. Cells classified as unassigned or showing ambiguous assignments were excluded from downstream analyses.

### Subtype refinement and iterative reannotation of the fetal atlas

To refine cell type annotations within the fetal atlas, specific neuronal and glial populations were subset from the integrated dataset and reanalysed. Dopaminergic and STN neurons, were first extracted based on harmonized annotations. Donor effects were corrected using BBKNN with DonorID as the batch variable after excluding donors represented by fewer than three cells. Leiden clusters (resolution = 1.8) were annotated based on canonical marker gene expression and assigned to dopaminergic and subthalamic lineages, including mDA neurons, A9 mDA neurons, mDA neuroblasts, STN neurons, STN neuroblasts, and a novel TH–PITX2 neuronal population. Hindbrain V2 neuronal populations were then further refined using a similar reanalysis strategy. Cells belonging to this lineage were reclustered using Leiden clustering (resolution = 0.8) and annotated into transcriptionally distinct subtypes based on marker gene expression, including glycinergic OTP-positive neurons, SOX14-positive glycinergic neurons, glutamatergic neurons and neuroblasts, GABAergic NR4A2-positive neurons, DTg populations, and progenitor states. Glial populations were curated separately from the Braun–Linnarson datasets. Cells corresponding to diencephalon, midbrain, hindbrain, pons, and cerebellum regions were retrieved. Glioblast lineage populations were reannotated based on Leiden clustering results (resolution = 0.1) and marker gene expression, assigning clusters to astrocytes or glioblasts. Similarly, oligodendrocyte lineage cells were annotated as oligodendrocytes or oligodendrocyte precursor cells (OPCs) according to cluster identity and canonical markers. Vascular populations were identified by selecting cells classified as vascular in the Braun dataset and were annotated using a CellTypist classifier trained on the human brain vascular atlas^135^, applying best-match prediction with majority voting. These refined annotations were then reconciled and concatenated back into the reference atlas to generate the final set of detailed fetal cell type labels. Integration was then done using CellHint for final visualization as this showed the best integration score (See: *Integration method comparison and benchmarking*).

### Differential expression analysis for fetal cell subtype signatures

To define fetal cell subtype signatures, differential expression analysis was restricted to genes annotated as protein-coding or long non-coding RNA (lncRNA) based on the gene_biotype annotation in the reference gene metadata. All other gene biotypes were excluded prior to analysis. To reduce imbalance across cell populations, up to 500 cells per detailed cell type were randomly sampled from the dataset for downstream analysis while preserving the overall cellular diversity of the atlas. Differential expression analysis was performed using Scanpy (scanpy.tl.rank_genes_groups) with the Wilcoxon rank-sum test, using Detailed_CellTypes as the grouping variable. Marker genes were retained if they passed an adjusted P value < 0.01 and had positive log fold change, indicating enrichment in the target subtype relative to other detailed cell types.

### Gene regulatory network inference

Gene regulatory networks for the mDA-STN and TH-PITX2 neurons were inferred using the pySCENIC pipeline^65^ following the standard three-step workflow of GRN inference, motif enrichment, and regulon activity scoring. Prior to analysis, genes were filtered to retain only those present in both the fetal expression dataset and the motif ranking feather files used for motif analysis. Gene regulatory network inference was performed using GRNBoost2 with a transcription factor list derived from the hg38 TF reference. In the second step, regulons were defined using the pySCENIC ctx module, which refines co-expression modules by motif enrichment analysis. Motif enrichment was evaluated using hg38 motif rankings (genes_vs_motifs.rankings.feather) together with the v10 motif annotation database (motifs-v10nr_clust-nr.hgnc). Modules were retained if the transcription factor–binding motif was significantly enriched among predicted target genes located near transcription start sites (TSS). Genes lacking direct TF-binding motifs were removed from the regulon. Only modules containing at least 20 target genes were retained, and motif enrichment was required to pass a normalized enrichment score (NES) threshold of 2.0. Dropout masking was applied during motif pruning (--mask_dropouts), and all potential modules were evaluated (--all_modules). In the final step, regulon activity in individual cells was quantified using the AUCell algorithm. Regulon activity profiles were subsequently used to compute Regulon Specificity Scores (RSS) for each cell population to identify lineage-specific regulatory programs.

### Spatial transcriptomics

For our Visium datasets, Visium Spatial Gene Expression for Fresh Frozen (10x Genomics) was performed following the manufacturer’s protocol. Cryosections of the frozen blocks were generated using a Leica CM1950 cryostat. Images were subsequently captured using a Hamamatsu S60 slide scanner at ×40 magnification before conducting the Visium protocol for subsequent alignment. Libraries were mapped with SpaceRanger (10X Genomics), to GRCh38-2020-A. Downstream analyses was performed with Scanpy. Count matrices were normalized by total counts and log-transformed using sc.pp.normalize_total and sc.pp.log1p, respectively. Highly variable genes were identified with the Seurat method (sc.pp.highly_variable_genes, flavor="Seurat") using the top 2,000 genes. To define spatial regions enriched for specific marker combinations, we performed a custom co-expression–based spatial mapping analysis. For each predefined marker set, expression values were extracted across all spatial spots. A gene-specific expression threshold was defined as the 40th percentile of expression across all spots, allowing thresholds to adapt to gene-specific expression distributions and reducing the influence of low background signal. Spots were considered positive for a given gene when expression exceeded this threshold. A spatial voxel was then classified as belonging to a given gene program only when all genes within the marker set simultaneously exceeded their respective thresholds, enabling the identification of areas defined by combinatorial marker expression rather than individual genes.

### Metadata curation and harmonization of mDA differentiation protocols for the HDCA

We compiled a collection of 15 published and 4 unpublished single-cell RNA sequencing datasets generated from human pluripotent stem cell–derived midbrain dopaminergic (mDA) differentiation protocols (Supplementary Table 1). For each dataset, raw count matrices and associated metadata were obtained from the corresponding publications, or directly from the authors for unpublished datasets. Metadata fields were harmonized across studies to generate a standardized annotation framework including the following variables: *Timepoint*, *Original annotation*, *Protocol*, *CellLine*, *Lab*, *Method*, and *Model*. Prior to integration, datasets were filtered to retain only relevant samples. Specifically, we excluded samples derived from disease models^2,13^ and treated with rotenone^8,13^. For assembloids only midbrain organoids were retained base on original cell type annotation^18,20^ . Finally, only timepoints of ≥14 days in culture were retained to ensure that floor plate patterning had been completed. Earlier timepoints were excluded because they may contain cell states not represented in our fetal reference atlas, such as pluripotent stem cells present at day 0. Remaining datasets were then concatenated into a single AnnData object using an outer join (join=’outer’) to preserve the union of detected genes across studies.

### Quality control and filtering of the HDNA atlas

Quality control filtering was performed on the integrated HDNA dataset using Scanpy. Putative doublets were identified using Scrublet^134^ with batch-aware processing based on the *Publication* metadata field, and predicted doublets were removed from downstream analyses. Cells expressing fewer than 200 genes were excluded, and genes detected in fewer than 5 cells were removed. Additional filtering was applied based on standard quality metrics. Cells with high mitochondrial transcript content (>15% of total counts), elevated ribosomal RNA content (>40%), or detectable hemoglobin gene expression (>0.05%) were excluded to remove low-quality cells. Cells with unusually high transcript counts (>40,000 total counts) or high number of genes (>6,000 detected genes) were also removed.

### Cell type classification of the HDNA using CellTypist

A neighborhood graph was constructed using 20 nearest neighbors and 30 principal components with a cosine distance metric, followed by UMAP embedding (*min_dist* = 0.1, *spread* = 3.5) for visualization. Batch effects across protocols and publications were corrected using BBKNN based on the *Protocol_Publication* annotation. Cell type prediction was performed using a fetal CellTypist classifier trained on the integrated fetal diencephalon–midbrain–hindbrain atlas without erythrocytes. Predictions were generated using two complementary modes: probability match, which assigns cell types based on probability thresholds (*p_thres* = 0.7) with majority voting across nearest neighbors, and best match, which assigns the most likely label for each cell.

### CapybaraBrain

CapybaraBrain decomposes each single-cell transcriptome into weighted contributions from predefined fetal reference programs, modeling cellular identity as a continuous mixture rather than enforcing discrete assignments. This formulation enables quantitative assessment of how closely a given cell aligns with its intended fetal program while simultaneously revealing competing or spurious programs within the same cell. Building on the conceptual framework of Capybara^26^, we adapted the mixture-weight inference and empirical significance testing approach into a marker-informed, scalable Python implementation tailored to our fetal atlas and the highly specific marker sets derived for each of the 93 fetal cell subtypes (Fig. 1, Extended Data Fig. 4).

#### Cell identity inference

For each of the 93 reference cell subtypes, a transcriptional program was defined as the mean normalized expression profile (the centroid) computed across all cells of that type in the fetal atlas, restricted to a selected marker gene set. The top 20 marker genes per cell subtype were used, selected based on specificity scores from differential expression analysis (see *Differential expression analysis for fetal cell subtype signatures*). The union of these marker genes across all 93 programs defined the feature space for inference.

For each query cell, expression values across this marker gene set were extracted from the normalized, log1p-transformed count matrix and square-root transformed prior to inference. This transformation was applied exclusively within the inference feature space, and the original expression matrix was preserved unchanged for all downstream analyses. Mixture weights were then estimated by finding the combination of reference centroids that best reconstructed the cell’s transformed expression profile, under the constraint that all weights were non-negative. This non-negativity constraint reflects the biological requirement that reference programs contribute additively to a cell’s transcriptional profile. After inference, weights were rescaled to sum to one to enable interpretation as proportional contributions. All steps, including centroid construction, transformation, and weight inference, were applied identically to both query and fetal reference cells, ensuring that downstream empirical significance testing was performed on a consistent scale.

#### Empirical significance testing of identity weights

To determine whether inferred weights reflect biologically meaningful identity assignments rather than background noise, we compared each cell’s weights against a null distribution derived from fetal reference cells. For each reference identity, the null distribution was built from fetal cells whose highest-weight identity was not that identity, capturing the distribution of incidental cross-identity weights expected in the absence of true assignment. For each cell and each identity, an empirical p-value was computed as the proportion of null weights equal to or exceeding the observed weight. To enable scalable null estimation, up to 60,000 fetal reference cells were used, subsampled with a fixed random seed when the reference exceeded this size.

#### Identity binarization and hybrid detection

Rescaled weights were converted to a binary identity matrix indicating which reference identities were significantly represented in each cell. An identity was retained when two criteria were jointly satisfied: the empirical p-value fell below a significance threshold and the rescaled weight exceeded a minimum contribution threshold. All results reported here used a significance threshold of α=0.05 and a minimum weight threshold of τ=0.15. Cells with no identity satisfying both criteria were labelled *unassigned*.

#### Hybrid-aware majority voting

To stabilize identity assignments and reduce noise from single-cell transcriptional variability, CapybaraBrain performs a hybrid-aware majority voting step across transcriptionally defined cell clusters. Highly variable genes were identified, expression values were scaled, and PCA was performed on the normalized expression matrix. A shared nearest-neighbour graph was constructed in PCA space and Leiden community detection was applied at high resolution to produce fine-grained over-clusters. Within each over-cluster, the most frequently assigned identity was determined and used to resolve ambiguous per-cell calls. Clusters in which no single identity reached a minimum proportion threshold were labelled *heterogeneous*. This cluster-level aggregation filters spurious assignments driven by transcriptional noise while preserving identity states that are consistently supported across transcriptionally similar cells.

For cells carrying multiple binarized identities, a hybrid-aware extension of this step was applied. Within each over-cluster, the most frequent identity pair (the dominant combination of primary and secondary identities) was determined. Cells whose secondary identity was inconsistent with the cluster-dominant pair were resolved to the primary identity alone, removing idiosyncratic multi-identity calls while retaining hybrid and transitioning states supported by cluster-level evidence.

#### Classification of discrete, hybrid and transitioning states

Cells retaining a single identity after majority voting were classified as *discrete*. Cells retaining two or more identities were further classified using a curated lineage hierarchy derived from the fetal atlas, which organizes cell states into developmental branches grouping progenitor, intermediate, and mature populations of the same lineage. Cells whose retained identities fell within the same branch were classified as *transitioning*, reflecting lineage-consistent progression along a developmental trajectory, for example mDA neuroblasts and mDA neurons within the midbrain dopaminergic lineage. Cells whose retained identities spanned distinct branches were classified as *hybrid*, reflecting partial engagement of transcriptional programs from developmentally unrelated lineages. Cells with no identity satisfying significance thresholds were labelled *unassigned* and clusters with no dominant identity were labelled *heterogeneous*.

### CapybaraBrain parameter benchmarking and comparison with CellTypist

To evaluate the robustness of CapybaraBrain identity assignments, we performed a systematic comparison of key model parameters, including the number of top reference programs retained during quadratic programming decomposition (TOPK), the empirical significance threshold used during identity binarization (α), and the minimum mixture weight required for identity retention (τ). For each parameter configuration, the full CapybaraBrain pipeline was executed through the transitioning-state detection step, generating identity annotations for both fetal reference cells and *in vitro*-derived cells. To assess classification performance, CapybaraBrain predictions were compared to annotations obtained using CellTypist with a custom classifier trained on the integrated fetal brain atlas (see *New human fetal sample processing and cell classification*). For each parameter configuration, datasets were down sampled to ensure a maximum of 200 cells per class. Agreement between CapybaraBrain and CellTypist predictions were calculated as the proportion of cells receiving identical identity assignments between majority vote calls (so not considering hybrids that CellTypist does not detect). For fetal cells, where curated reference annotations were available, additional performance metrics were computed using the reference labels as ground truth. These included overall classification accuracy as well as macro-averaged and weighted precision, recall and F1 scores for both CapybaraBrain and CellTypist predictions. Performance metrics were summarized across parameter combinations to identify parameter regimes that maximized agreement with reference annotations and overall classification performance. CellTypist achieved a mean accuracy of 91.28% and a macro-F1 of 0.901. Using the evaluated CapybaraBrain configuration (TOPK = 20, α = 0.005, minimum weight = 0.15), CapybaraBrain achieved a mean accuracy of 92.53% and a macro-F1 of 0.902, representing an improvement of 1.25 percentage points in accuracy and 0.001 in macro-F1 over CellTypist (Extended Data Fig. 7c). To obtain a conservative assignment of A9 dopaminergic neurons, as this subtype has very few markers that distinguish it and is an important target of many protocols, cells predicted as *MB A9 mDA neurons* by the CapybaraBrain majority-voting label were retained as A9 only when their transferred CellTypist fetal best-match annotation also corresponded to *MB A9 mDA neurons*; otherwise they were reassigned to the broader *MB mDA neurons* class. This additional constraint was introduced because A9 identity is defined by a small number of highly specific markers, such as *ALDH1A1*, that distinguish this subtype from closely related dopaminergic populations, while many other markers of A9 are related to maturation rather than identity (like *SLC6A3*). While the CapybaraBrain framework evaluates similarity across gene programs, the CellTypist classifier applies gene-specific weights learned from reference data, enabling stronger emphasis on subtype-defining markers and we observed improved discrimination between highly related dopaminergic states. Using these assigned cell identities, we performed label-aware data integration and systematically compared multiple integration strategies within a previously established benchmarking framework^136^. Among the tested approaches, CellHint^56^ demonstrated the best overall performance for integrating these datasets (Extended data Fig. 7b).

### Harmonization of adult post-mortem midbrain dopaminergic neuron subtypes

Single-cell datasets from the adult human brain studies by Siletti^53^ and Kamath^52^ were integrated and their midbrain dopaminergic (mDA) subtype annotations harmonized using CellHint. Highly variable genes were identified separately within each dataset using scanpy.pp.highly_variable_genes, specifying the dataset of origin as the batch_key. Cell type alignment across datasets was then performed using cellhint.harmonize, using dataset identity as the batch variable and the original mDA subtype annotations from each study as input labels. Manual curation was subsequently performed using study-specific marker genes to establish a unified nomenclature for each dopaminergic subtype across datasets. Guided integration was then performed using CellHint, using the integrated AnnData object as input and specifying DonorID as the batch variable and the curated Harmonized_CellType annotations as reference labels (n_meta_neighbors = 1)

### Integration method comparison and benchmarking for fetal, adult and HDNA datasets

To evaluate the performance of different batch-correction and data integration strategies, we benchmarked multiple integration methods across fetal, adult, and *in vitro* dopaminergic neuron datasets using a unified evaluation framework. Precomputed embeddings generated by several commonly used integration approaches were collected and compared, including unintegrated data (baseline), Harmony^137^, Scanorama, scVI^138^, scANVI^139^, CellHint^56^ (default and guided), LIGER^140^, BBKNN^141^, ComBat^142^, and STACAS^143^. Integration performance was assessed using the scIB benchmarking framework^136^, which evaluates both batch correction and biological conservation. Batch correction performance was quantified using metrics including Batch ASW (BASW), iLISI, graph connectivity, and principal component regression comparison, while biological conservation was evaluated using label-based metrics that assess preservation of cell type structure after integration. For the fetal datasets, benchmarking was performed using DonorID as the batch variable and Detailed_CellTypes as the biological label reference. For adult midbrain dopaminergic neuron subtypes, the same framework was applied using DonorID as the batch variable and Harmonized_Labels as the reference annotation. For the *in vitro* HDNA datasets, Publication was used as the batch variable and cell type labels assigned using Capybara majority voting were used as the biological reference. For each dataset and integration method, the corresponding embedding was evaluated using the Benchmarker class implemented in scib-metrics, combining both BatchCorrection and BioConservation evaluation modules.

### Fetal and adult midbrain dopaminergic neurons maturation GRNs and gene modules

To identify modules associated with maturation of mDA neurons, we restricted our analysis to single-nuclei RNA-seq data of MB A9 mDA neurons in fetal cells and SOX6_AGTR1 in adult post-mortem tissue of age 50 years old or below. The two datasets were concatenated and then gene expression matrices were normalized per cell and log-transformed. Highly variable genes were identified in a batch-aware manner across datasets. Neighbourhood graph was then constructed using 50 nearest neighbours in PCA space with cosine distance, and a two-dimensional embedding was generated using UMAP. Prior to module discovery, analyses were restricted to genes annotated as protein-coding or long non-coding RNAs. Hotspot^144^ was used to find gene modules. A neighbourhood graph with 30 nearest neighbours was used to compute gene-wise autocorrelation. Genes with a false discovery rate (FDR) < 0.05 were considered significantly autocorrelated. The top 500 genes ranked by autocorrelation Z-score were retained for downstream analysis, and pairwise local gene–gene correlations were computed to define co-expression modules. To identify GRNs the same pipeline described in “*Gene regulatory network inference*” using the same fetal and adult subset. Gene ontology (GO) enrichment analysis was done using the GO Biological Process gene set collection. Enriched terms were identified for each module and ranked according to adjusted P values. Only terms with FDR ≤ 0.05 were considered significant. To focus specifically on neuronal maturation programs, enriched GO terms were filtered using predefined keyword categories representing axonogenesis and neurite development (e.g., *axonogenesis, axon guidance, neurite outgrowth, neuronal projection, growth cone, dendrite*), synaptic processes (e.g., *synapse, synaptic signalling, neurotransmitter, presynaptic, postsynaptic, vesicle, exocytosis*), and neuronal differentiation (e.g., *neurogenesis, neuron differentiation, neuronal fate*) (Supplementary Table 6). To visualize relationships between enriched biological processes and their associated genes, a gene–ontology network was built linking enriched GO terms to module genes. For visualization clarity, up to eight enriched GO terms per module and eight associated genes per term were retained. Network layouts were generated using a force-directed graph algorithm.

### Module-based fidelity scoring against fetal and adult mDA modules and TFs

To quantify the transcriptional similarity of each *in vitro* model to *in vivo* references, we computed correlation-based fidelity scores at the level of timepoint–protocol–publication (TPP) groups. Gene module scores were calculated using Scanpy’s score_genes function across ten co-expression modules and SCENIC enriched TFs derived from fetal and adult mDA neurons. To account for systematic technical differences between single-cell and single-nucleus RNA-seq measurements, we projected each cell’s module score vector onto the SC–SN bias direction, defined as the mean score difference between modalities, normalized to unit length, and removed this component via linear regression, re-centering to the global mean. Tech-corrected scores were then averaged per TPP group (minimum 10 cells), and Pearson correlations were computed between each TPP mean profile and the mean profiles of fetal and adult reference populations, yielding two fidelity coordinates per TPP: correlation with fetal and correlation with adult. TPP groups were assigned to panels (monolayer, organoid, graft, or reference) based on the model annotation. *In vitro* reference anchors were defined as the mutual correlation between fetal and adult mean profiles, plotted as fixed landmarks in all panels to enable cross-panel comparison.

### Glycolysis, hypoxia, ER stress, mDA fidelity and maturation scoring across *in vitro* and *in vivo* dopaminergic models

Cells were scored for glycolysis, hypoxia, ER stress, mDA fidelity and neuronal maturation using Scanpy score_genes. Glycolysis and hypoxia signatures were defined using MSigDB Hallmark gene sets, ER stress using the Gene Ontology term *response to endoplasmic reticulum stress*, mDA fidelity was scored on the top 50 marker genes for fetal MB mDA neurons, maturation was scored using Module 8 specifically that was enriched in adult only mDA neurons combined with adult specific SCENIC-derived TFs. To account for systematic differences between single-cell and single-nucleus measurements, gene set scores were corrected independently within each modality. Scores were first centered by subtracting the modality-specific mean and were then re-scaled to the global mean across all cells, thereby removing modality-driven offsets while maintaining the relative variation in scores within each modality. Corrected scores were summarized across grouped timepoint–protocol–publication categories, excluding groups with fewer than 10 cells.

### Composite mDA identity and maturation scoring across *in vitro* and *in vivo* models

To summarize overall *in vitro* model performance, we derived a composite score integrating mDA identity and neuronal maturation. For each TPP group (minimum 10 cells), median tech-corrected scores were computed for mDA identity (top 50 fetal MB mDA marker genes) and neuronal maturation (Module 8). Each score was independently z-scored across TPPs and summed to yield a single composite fidelity metric per TPP.

### Within-protocol/publication, time-controlled correlations between metabolic and stress-related programs and neuronal states

To control for protocol- and publication-specific baseline effects, scores were mean-centered within each protocol–publication group. Residual variation not explained by time in culture was then obtained by regressing these values against time using a second-degree polynomial model. Then spearman correlations were computed between glycolysis, hypoxia and ER stress scores and mDA identity or maturation scores using these within-protocol/publication, time-corrected residuals.

### Nearest-neighbor projection of PD tricultures onto the HDNA atlas

To enable mapping of the PD triculture query dataset onto the Human Dopaminergic Neuron Atlas (HDNA), cells from the Studer triculture dataset contained within the reference atlas were split in a stratified 50:50 manner based on their CAPYMV_VITRO_majority_voting cell-type annotations (random seed = 7). One half of these cells was retained in the reference atlas, while the held-out half was added to the query object as wild-type internal controls, allowing downstream assessment of mapping fidelity. The combined query object was normalised to 10,000 counts per cell and log1p-transformed. For mapping, query data were restricted to the intersection of the query variable genes and the highly variable genes defined in the reference atlas. Query cells were then projected into the reference PCA space by matrix multiplication against the reference loading matrix (50 principal components), without re-fitting PCA on the query data. Nearest-neighbour mapping was performed using a k-nearest neighbour search in PCA space (Euclidean distance, k = 50; scikit-learn NearestNeighbors). For each query cell, the mean and minimum Euclidean distances to its 50 nearest reference neighbours were recorded as quantitative measures of atlas proximity. Cell-type label transfer was performed by majority voting across the k nearest neighbours independently for three annotation levels: *in vitro* majority-voting labels (CAPYMV_VITRO_majority_voting), hybrid-aware final annotations (CAPYMV_VITRO_hybrid_mv_final_call), and fetal best-match classifications (Fetal_CellClassification_BestMatch_majority_voting). The confidence of each assignment was defined as the fraction of the k neighbours supporting the predicted label. To generate a stricter assignment for A9 dopaminergic neurons, cells predicted as MB A9 mDA neurons by majority voting were retained in this class only if their independently transferred fetal best-match label also resolved to MB A9 mDA neurons; otherwise they were downgraded to the broader MB mDA neurons category. Cells with a label-transfer confidence below 0.4 were flagged as low-confidence and excluded from downstream analyses. For visualisation, mapped cells were projected onto the atlas UMAP by assigning each query cell the UMAP coordinates of its single closest reference.

### Data availability

Data will be available upon publication.

### Code availability

CapybaraBrain code is available at https://github.com/VittoriaDBocchi/CapybaraBrain

## Acknowledgements

This publication is part of the Human Cell Atlas –www.humancellatlas.org/publications/. We thank members of the Studer, Teichmann, Betel, Parmar, Kunath, Croft and Tabar lab for discussions on the manuscript. We would also like to thank the Integrated Genomic Operation (IGO) and the Single Cell Analytics Innovation Laboratory (SAIL) at MSKCC for outstanding technical support. Cores are supported by the NCI Cancer Center Support Grant (P30 CA08748), Cycle for Survival, and the Marie-Josée and Henry R. Kravis Center for Molecular Oncology. The work was supported in part by NIH grants 1R01NS118067 and 2R01AG054720 (L.S., D.B.), 1R01NS128087 (L.S.), R01NS126588 (V.T.), support by the National Research Foundation of Korea (NRF) grant funded by the Korea government (MSIT) (No. RS-2024-00351442) and the POSCO Science Fellowship of POSCO TJ Park Foundation to T.W.K., Support from the Freedom together Foundation (FtF) to L.S., and from the Marie-Josée Kravis Women in Science Endeavor and Aligning Science Across Parkinson’s (ASAP) to V.D.B. This work was also supported by funding to M.P. from Swedish Research Council (2021-00661), Swedish Parkinson Foundation (Parkinsonfonden), Swedish Brain Foundation. R.A. was funded by Aligning Science Across Parkinson’s (ASAP-020600) through the Michael J. Fox Foundation for Parkinson’s Research (MJFF), and NIH-R01NS119690-01. T.K. was supported by AMED Grant Number JP20bm0704054 and MRC Regenerative Medicine Grant Mr/V00560x/1 and by AMED under Grant Number JP20bm0704054 to A.M. ASAP-000472 also supported J.E.P., G.F.C. This work was also supported by the DFG (Project-ID 388169927 to the Microscope Core); the Alexander von Humboldt Foundation (Humboldt Research Fellowship for Experienced Researchers) and the BONFOR program (2024-1B-04) of the Medical Faculty, University of Bonn to Y.S.-I., and the iBehave project funded from the program ‘Netzwerke 2021’, an initiative of the Ministry of Culture and Science of the State of North Rhine-Westphalia to S.B..

## Author contributions

Conceptualization: V.D.B. and L.S.

Computational analysis and interpretation of the data: V.D.B. with contributions from K.T., P.Z., P.S., J.H., C.X., L.S., S.A.T.

Human fetal sample collection and droplet and spatial data generation and quality control: K.T., F.M., C.X., O.A.B., X.H., R.A.B. and S.A.T.

Methodology and development of CapybaraBrain: V.D.B.

Dataset integration and benchmarking: P.Z. and D.B.

Tri-culture differentiation of control and Parkinson’s disease lines and single-cell library preparation: L.W. Boost and Boost+ neuron differentiation and *in vivo* transplantation: D.Y., T.W.K., S.Y.K., J.P., V.T. and L.S.

Differentiation, single-cell preparation, alignment and quality control of assembloids and monolayer Parmar datasets: P.S., A.F., E.S., and M.P.

Differentiation, single-cell preparation, alignment and quality control of monolayer Kunath datasets: N.J.D., N.B.C., A.C., A.M., and T.K. Supported by AMED Grant Number JP20bm0704054 and MRC Regenerative Medicine Grant Mr/V00560x/1 and by AMED under Grant Number JP20bm0704054 to A.M.

Differentiation, single-cell preparation, alignment and quality control of organoid Croft datasets: K.K.S., J.P., J.Pow, and G.C.

Mouse TH–PITX2 validation experiments: L.K. and R.A. L.K. performed and analysed RNAScope based identification of Th-Pitx2 neurons in mouse brains, with supervision from R.A.

Mouse genetic fate-mapping experiments: Y.S-I. and S.B. Designed and performed genetic inducible fate-mapping experiments, performed immunostaining, acquired and analysed imaging data.

Supervision: L.S. and S.A.T.

Funding acquisition: L.S., S.A.T., D.B.

Writing – original draft: V.D.B., K.T., S.T. and L.S. Writing – review and editing: all authors.

---

**Extended Data Fig. 1.**
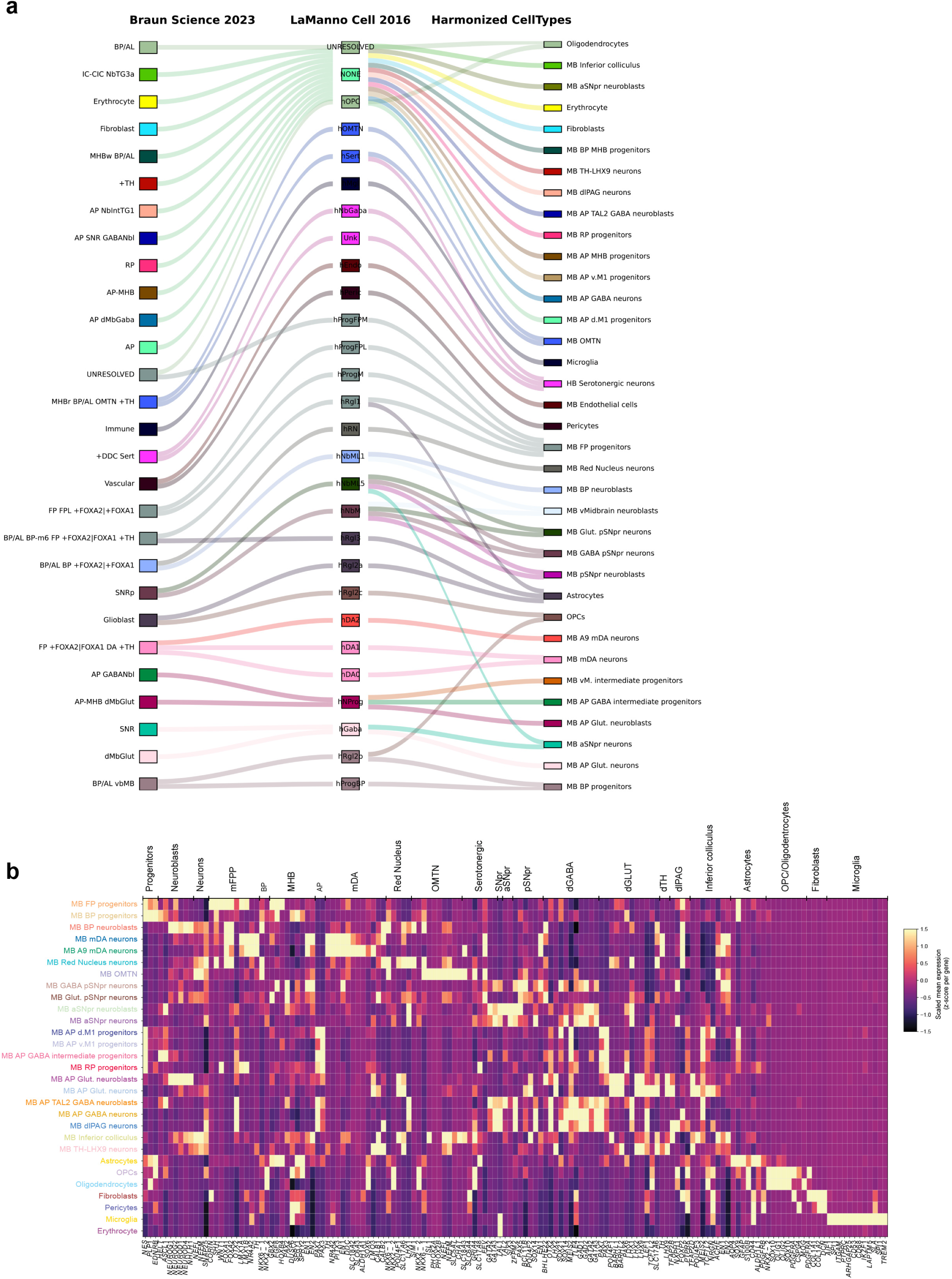
Label reconciliation and manual curation of fetal midbrain cell types. **a**, Sankey plot showing CellHint alignments between the Braun *et al*. Science, 2023 and La Manno *et al*. Cell, 2016 datasets and the final manually curated harmonized cell types. CellHint labels unmatched cell types as “NONE”, and cells that fail to integrate into the harmonization graph after the final iteration as “UNRESOLVED”. These unresolved cells are primarily from the Braun dataset, which includes dorsal midbrain regions not sampled by La Manno *et al*. Key differences were identified across studies; for example, in the Braun dataset, midbrain dopaminergic (mDA) neurons were broadly annotated as FP+FOXA1|FOXA2 DA+TH cells, whereas the La Manno study resolved these cells into three subtypes (DA0, DA1 and DA2). Based on the A9 mDA specific marker *ALDH1A1*, we unified these annotations by defining DA0 and DA1 as broad mDA neurons and DA2 as A9 mDA neurons, we did not find strong evidence of A10 neurons at this stage. **b**, Heatmap of expression values of canonical marker genes used for manual curation of midbrain cell subtypes.

**Extended Data Fig. 2.**
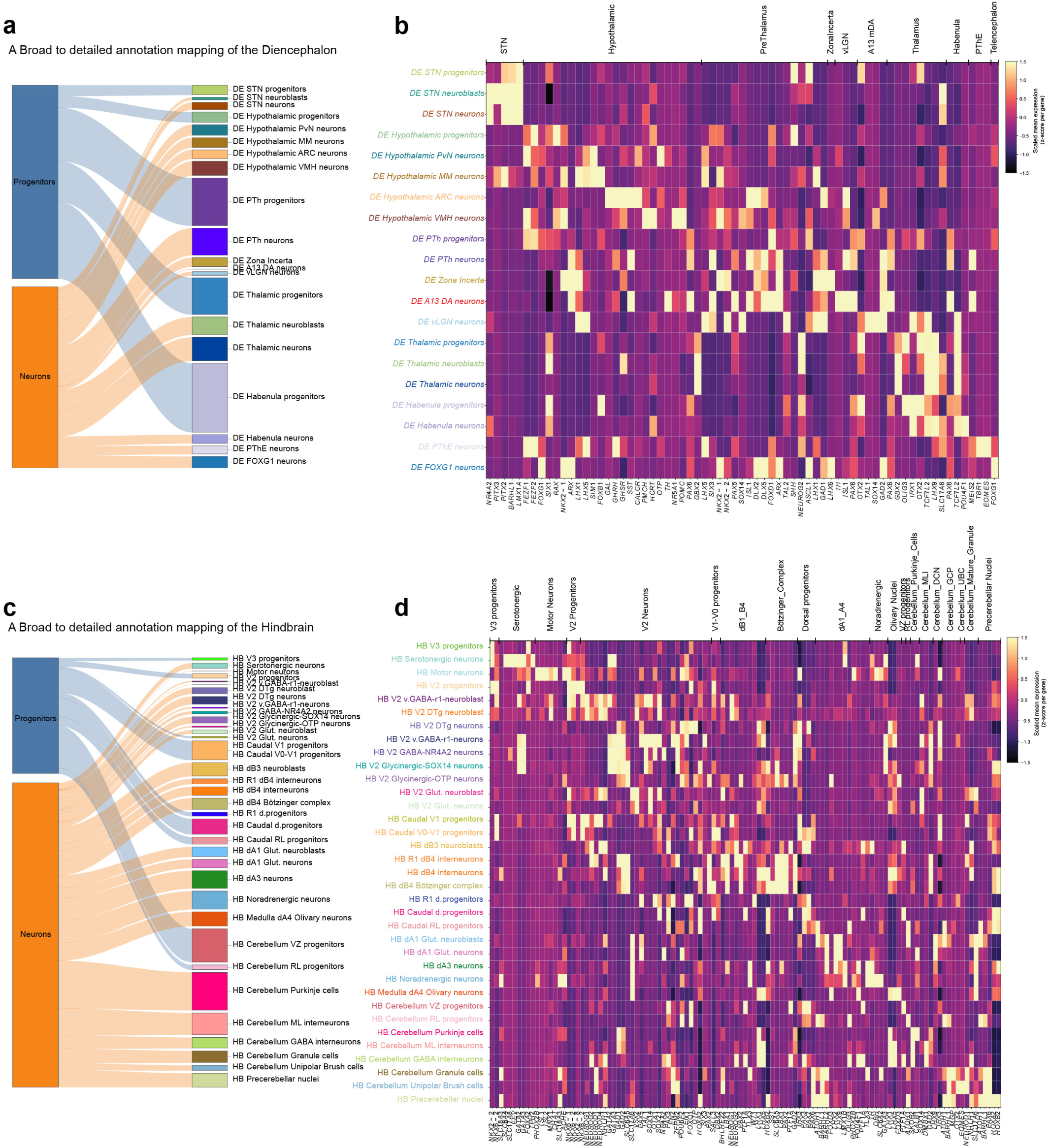
Detailed cell-type annotation of the fetal diencephalon and hindbrain. **a**, Sankey plot showing reannotation of cell subtypes from Human Cell Atlas Development (v1) in the diencephalon to refined cell-type annotations (v2). **b**, Heatmap of canonical marker genes used for manual curation of diencephalic cell states. **c**, Sankey plot showing reannotation of cell subtypes from Human Cell Atlas Development (v1) in the hindbrain to refined cell-type annotations (v2). **d**, Heatmap of canonical marker genes used for manual curation of hindbrain cell states.

**Extended Data Fig. 3.**
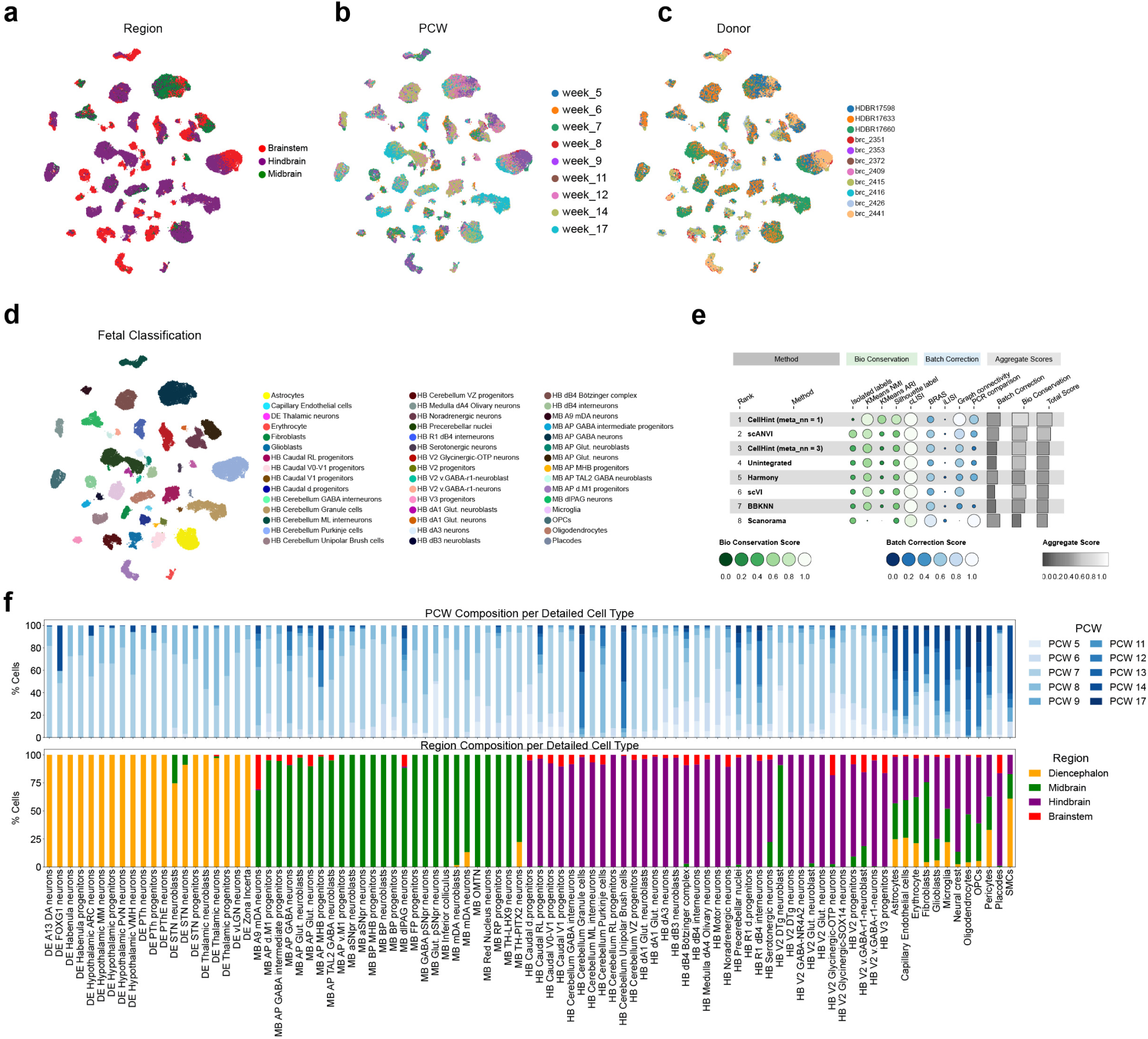
Classification and integration of newly generated fetal brainstem samples by single-nucleus RNA sequencing. **a-d**, UMAP coloured by sample origin (midbrain, hindbrain or whole brainstem) (a), PCW (b), donor identity (c), and CellTypist classification using the integrated atlas with v2 detailed annotations (d). **e**, Comparative benchmarking of data integration methods for incorporating newly generated datasets into the Braun et al. Science 2023 atlas, assessed using multiple quantitative metrics capturing batch correction and biological conservation. **f**, Stacked bar plot showing PCW and regional composition across detailed cell types.

**Extended Data Fig. 4.**
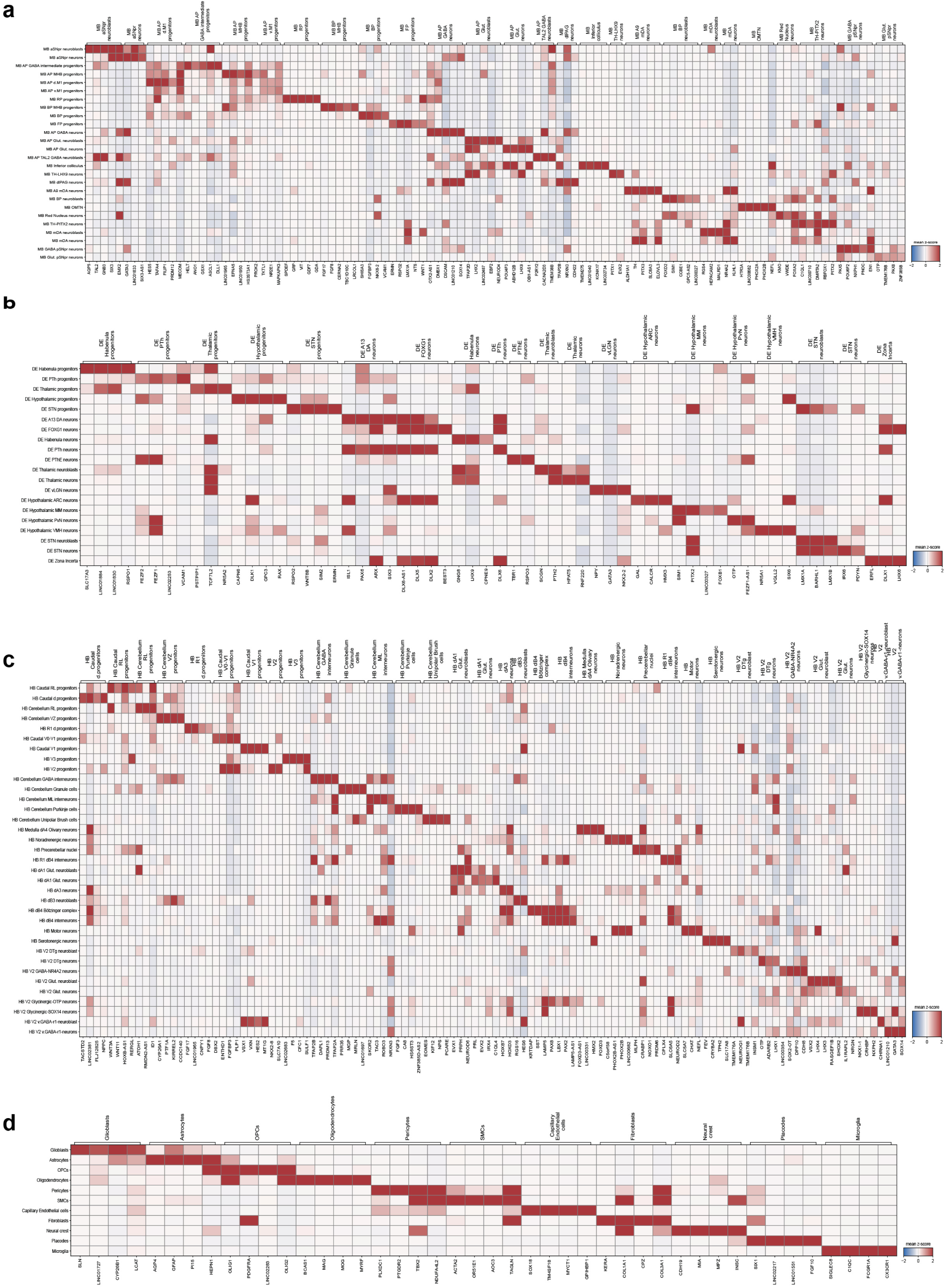
Marker gene dictionary across the fetal caudal brain. **a-d**, Heat maps showing cell-state-specific marker genes of the 93 cell states divided in midbrain (a), diencephalon (b), hindbrain (c) regions, and non-neuronal populations including glial cells, mural cells and fibroblasts (d).

**Extended Data Fig. 5.**
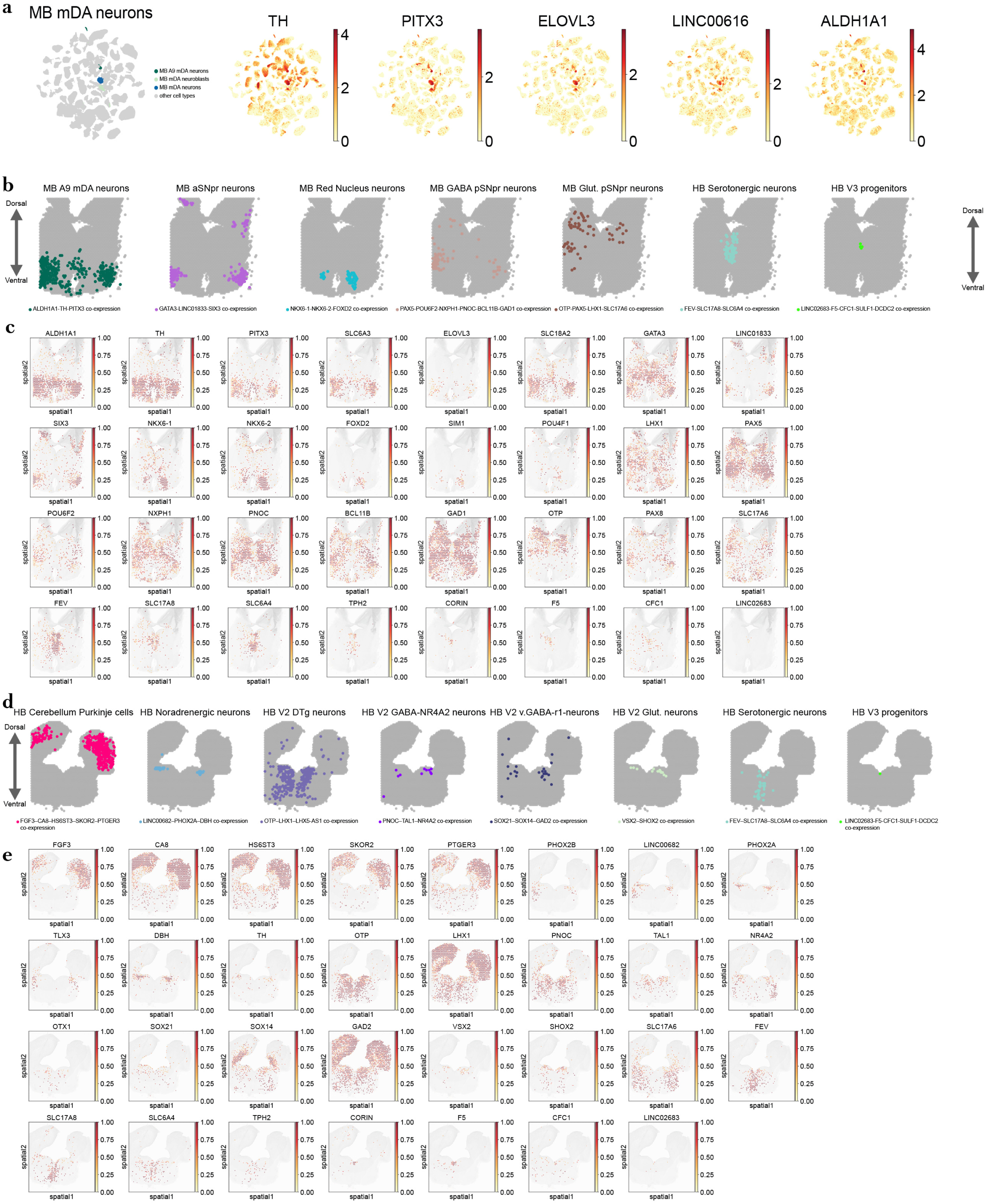
Spatial validation of ventral midbrain and hindbrain cell states. **a**, UMAP of highly specific marker genes of mDA and A9 mDA neurons. **b-c**, Spatial co-expression of key marker gene combinations defining ventral midbrain neuronal populations, including anterior SNpr neurons, red nucleus neurons, GABAergic and glutamatergic posterior SNpr neurons, and A9 mDA neurons. **d-e**, Spatially resolved heat maps across a 10 PCW hindbrain section showing canonical and newly identified cell states and marker genes. Top panels show co-expression of defining marker genes in each voxel (b,d), and bottom panels show spatially resolved expression heat maps of canonical and newly identified marker genes across 10 PCW sections (c,e).

**Extended Data Fig. 6.**
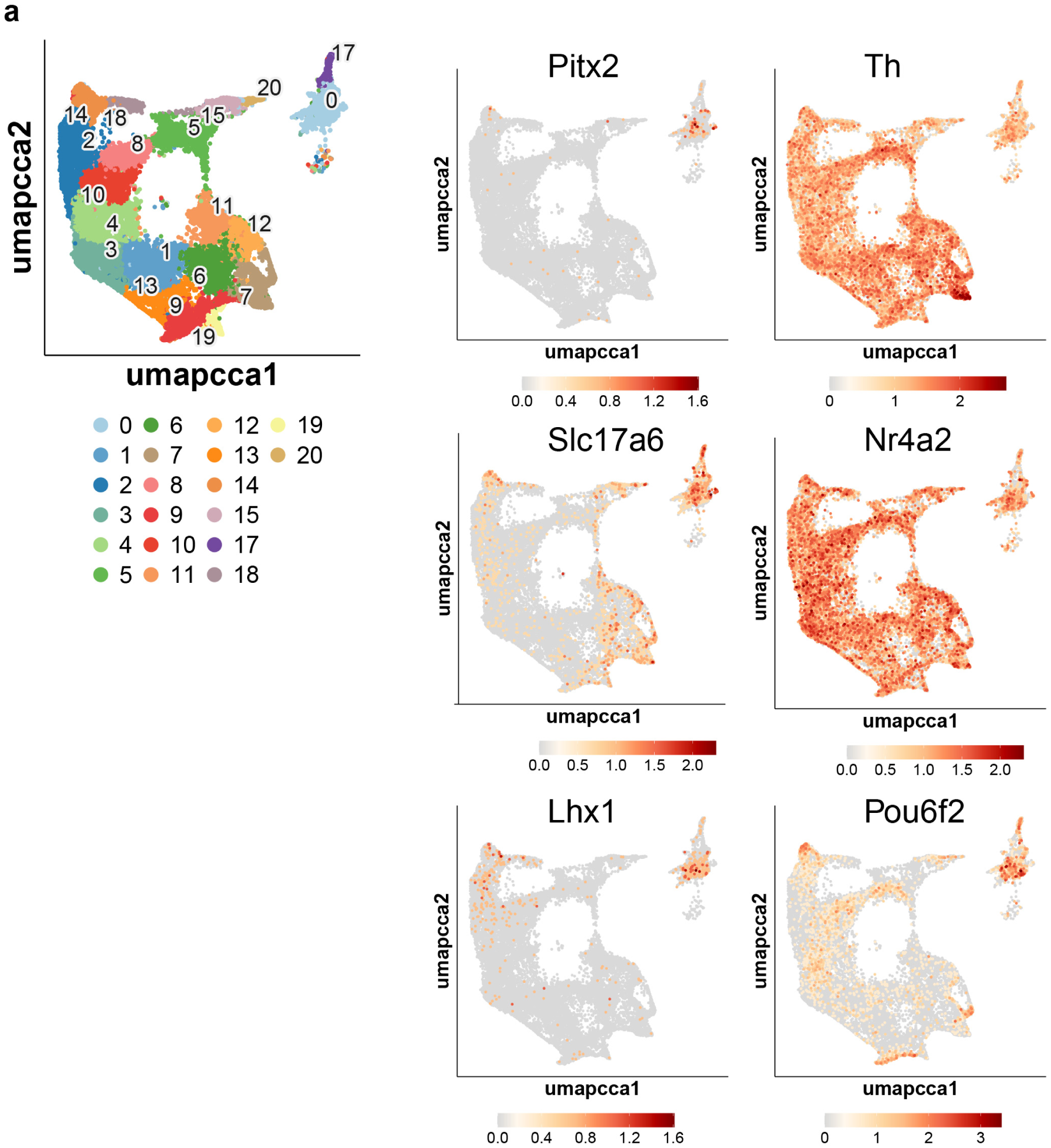
snRNA-seq from dopaminergic nuclei of the mouse brain. **a,** UMAP of snRNA-seq from published dopaminergic nuclei (defined by Dat-Cre expression) of 6-month old mice showing cluster 0 co-expressing *Pitx2*, *Th*, *Slc17a6*, *Nr4a2*, *Lhx1* and *Pou6f2*.

**Extended Data Fig. 7.**
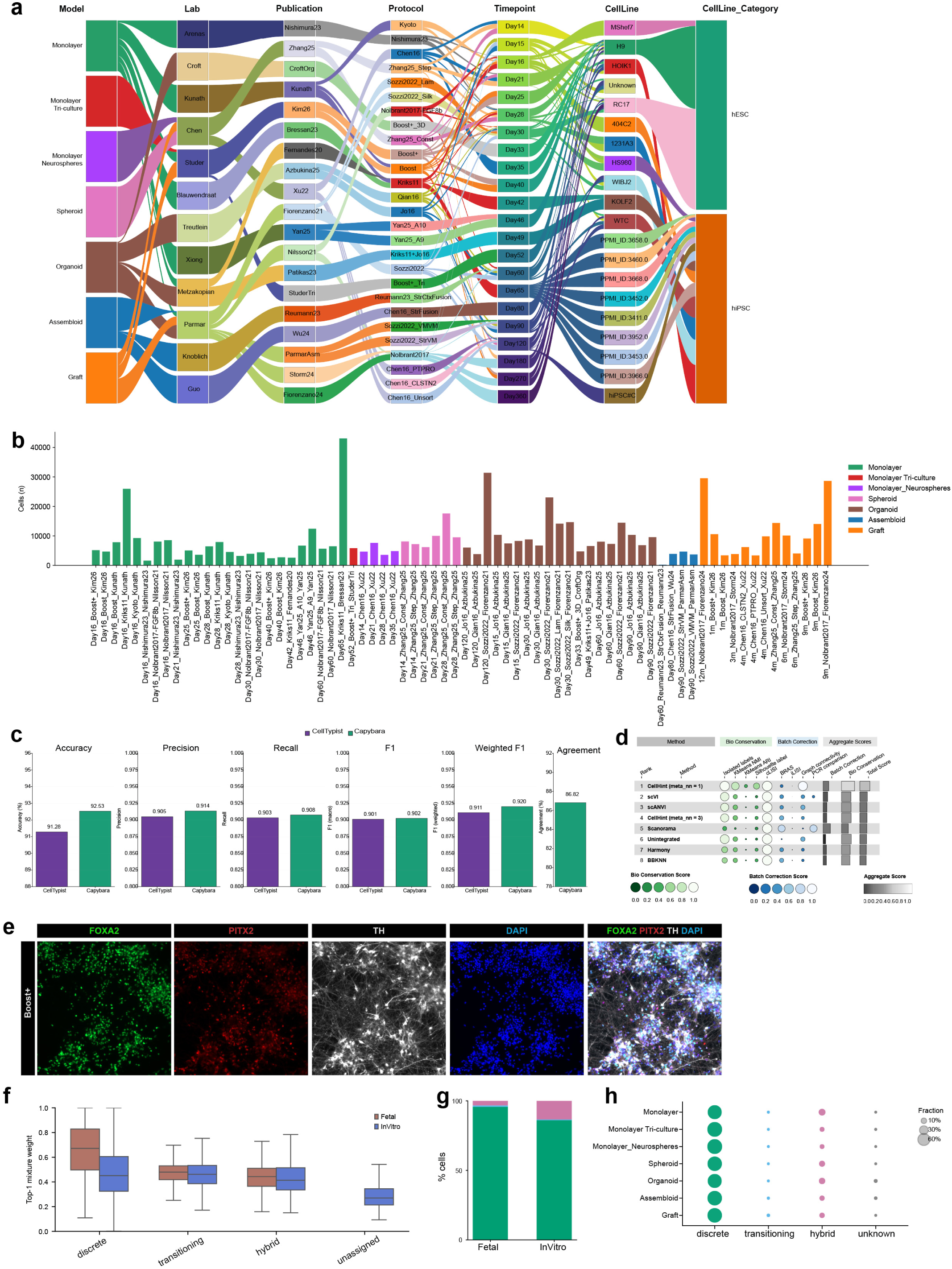
In vitro datasets integration and benchmarking of CapybaraBrain. **a,** Sankey plot summarizing datasets integrated into the HDNA, showing relationships between models, protocols, publications, laboratory that generated the data, timepoints and cell lines. **b,** Bar plots summarizing the number of cells contributed by each timepoint, protocol, and publication. **c,** Performance benchmarking of CapybaraBrain compared to CellTypist using the fetal reference atlas with ground-truth labels. Bars show accuracy, macro-precision, macro-recall, macro-F1, weighted-F1, and agreement between CapybaraBrain and CellTypist predictions. **d,** Comparative benchmarking of integration methods for harmonizing all *in vitro*-derived datasets. Methods were evaluated using quantitative metrics assessing batch correction and biological conservation. **e**, Immuno-fluorescent staining for TH-PITX2 neuron markers: TH, PITX2, FOXA2 and DAPI at day40 *via* the Boost+ conditions. **f**, Box plots show the distribution of Top-1 mixture weights across discrete, transitioning, hybrid, unassigned, unknown states. In fetal tissue, discrete cell states exhibited the highest Top-1 weights, consistent with well-resolved and transcriptionally dominant lineage identities. Hybrid and transitioning states displayed intermediate scores in both datasets, as expected for mixed or intermediate states. The “unknown” category was observed exclusively *in vitro*, likely representing populations not captured by the fetal reference or cells lacking a clearly dominant lineage program. **g**, Stacked bar plot showing the fraction of cells assigned to each CapybaraBrain state category (discrete, transitioning, hybrid, unknown) in fetal and *in vitro* datasets. **h**, Dot plot showing the fraction of cells in each CAPY_state_step4 category across *in vitro* model classes.

**Extended Data Fig. 8.**
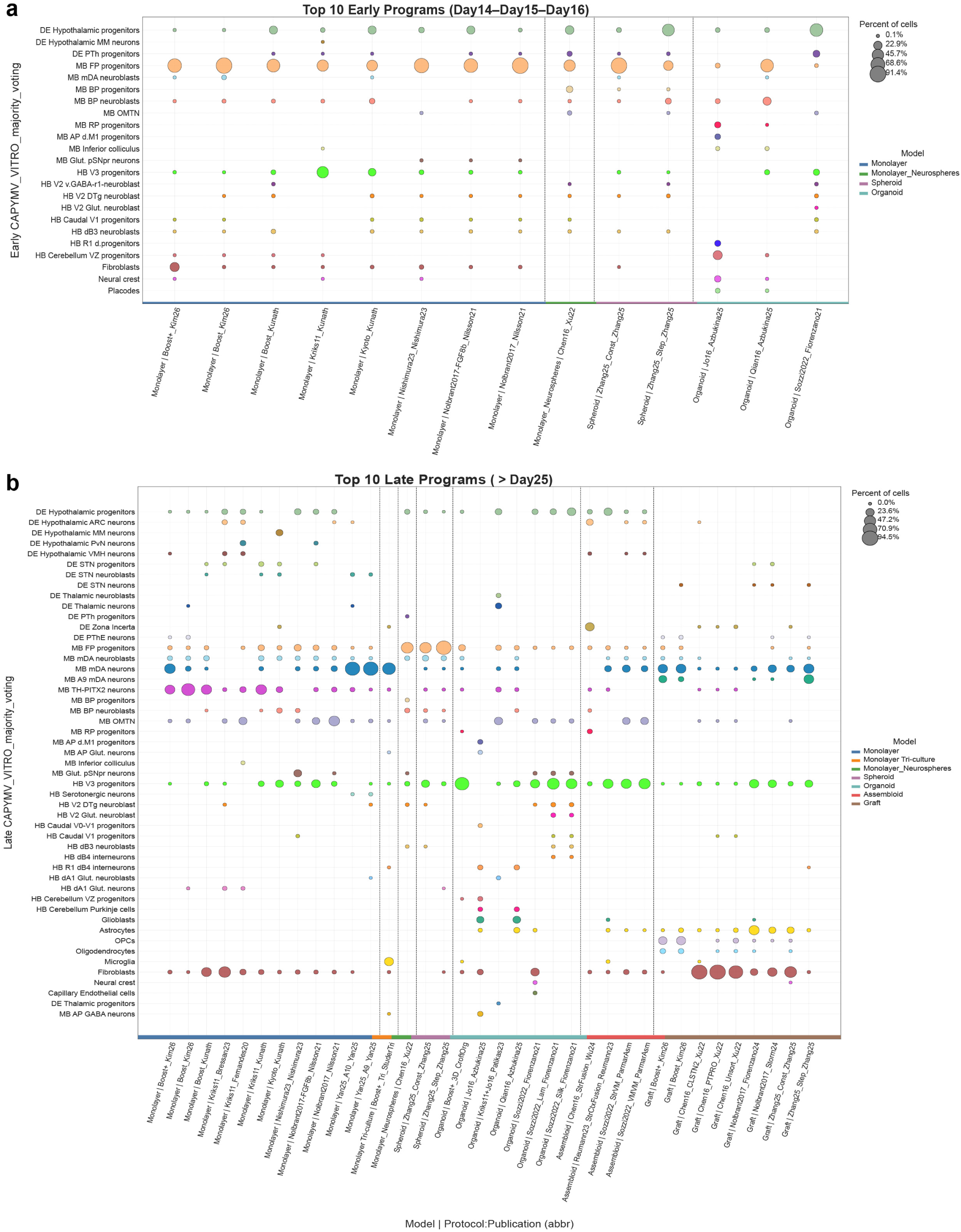
Cell-type composition across early and late differentiation timepoints. **a-b**, Bubble plot showing the top 10 most abundant cell types within early timepoints (Day 14–16) for each model/protocol/publication (a) and after Day ≥ 25 (b).

**Extended Data Fig. 9.**
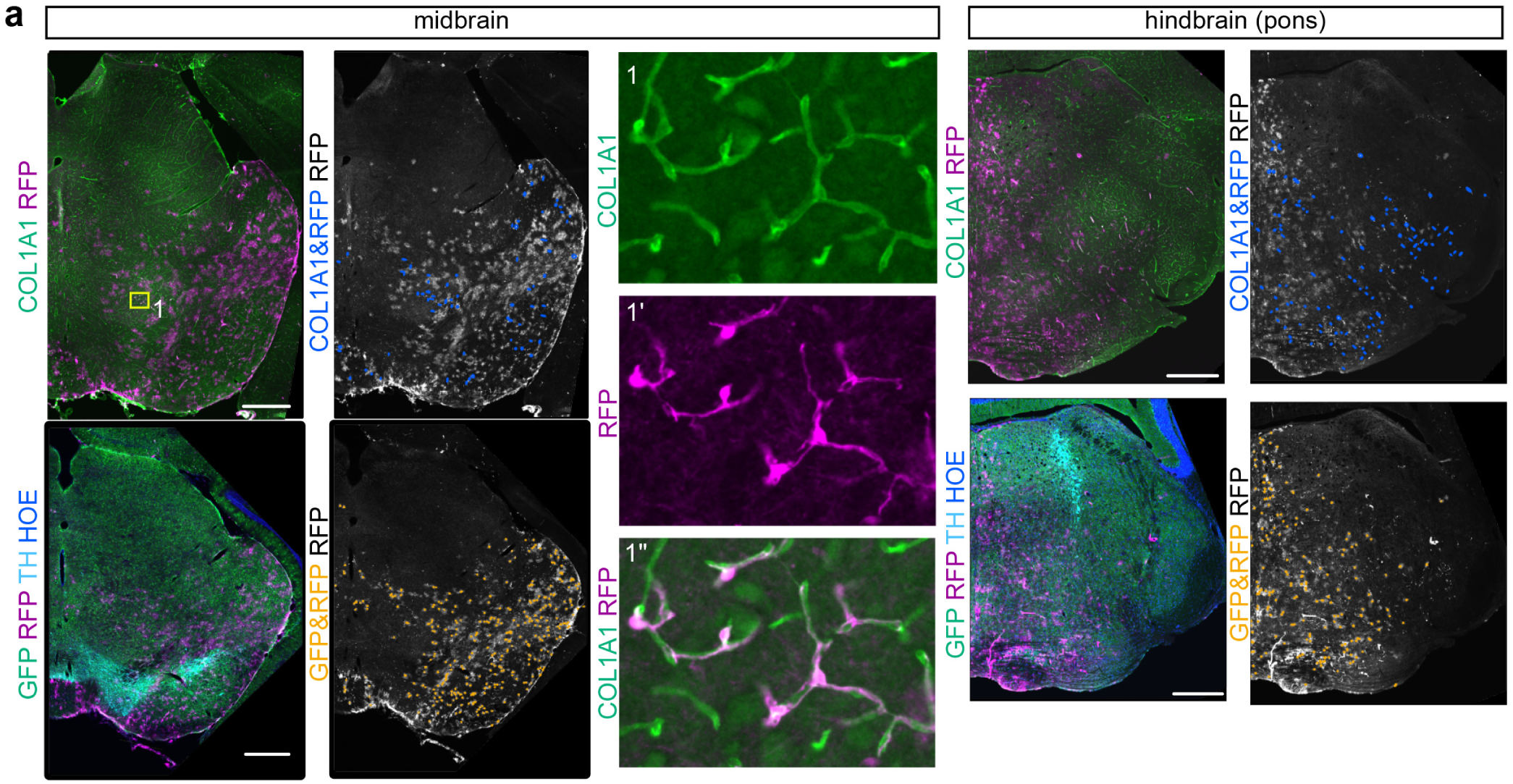
Fate mapping of Gli1-expressing progenitors in the mouse midbrain and hindbrain. **a**, Coronal sections of the midbrain and hindbrain of a *Gli1^CreER/+^, R26^Ai9/+^, Aldh1l1-GFP* mouse treated with TM at E9.5. Upper panels: Immunostaining for COL1A1 (fibroblasts) and RFP (fate-mapped cells) (left panels). Cells co-expressing COL1A1 and RFP are indicated by blue dots (right panels). (1–1″) Higher-magnification images of boxed regions highlight representative COL1A1+/RFP+ cells. Bottom panels: Immunostaining for GFP (ALDH1L1+ astrocytes), RFP (fate-mapped cells) and tyrosine hydroxylase (TH, monoaminergic neurons) (left panels). HOE: Hoechst. TH labels dopaminergic neurons in the midbrain and noradrenergic neurons in the pons. Cells co-expressing GFP and RFP are indicated by orange dots (right panels). The distribution of COL1A1+/RFP+ cells is comparable to that of ALDH1L1+ astrocytes co-expressing RFP and GFP suggesting that *Gli1*-expressing progenitors in the ventral mid- and hindbrain may give rise to both fibroblasts and astrocytes. Three mice were evaluated and similar distribution was confirmed in all mice.

**Extended Data Fig. 10.**
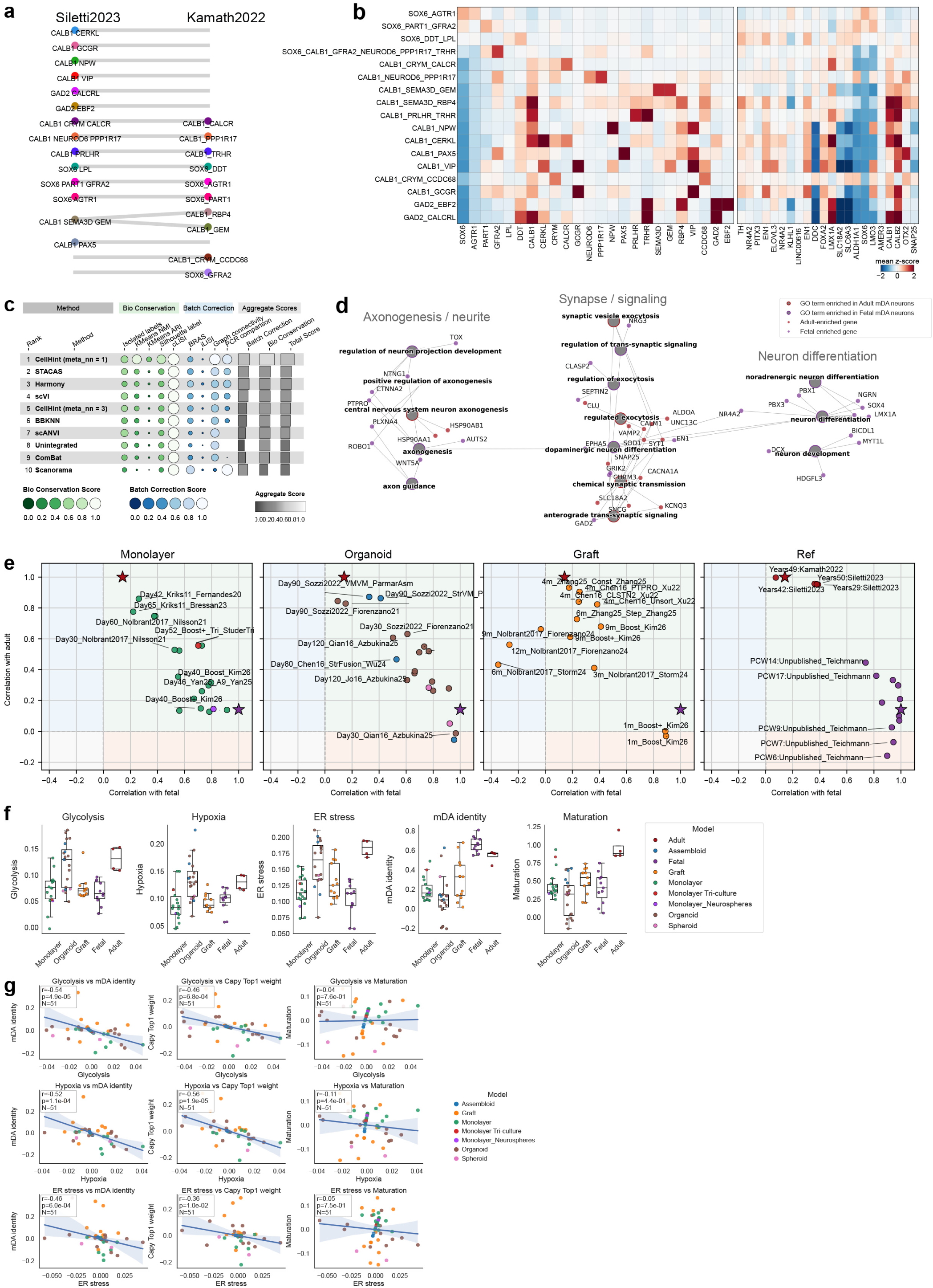
An integrated atlas of adult mDA neurons subtypes and maturation linked modules. **a**, Harmonization graph showing cell-type relationships between mDA subtypes in the Kamath study and the Siletti study. The Siletti atlas identified several subtypes that were not reported in the Kamath dataset. Several equivalent populations were labelled differently across studies. For instance, the SOX6-LPL neurons described in Siletti correspond to the SOX6–DDT population reported in Kamath, indicating that these represent the same underlying cell state despite differences in nomenclature. **b**, UMAP coloured by dataset publication **c**, Comparative benchmarking of data integration between the Kamath study and the Siletti study, assessed using multiple quantitative metrics capturing batch correction and biological conservation. **d**, Network representation of significantly enriched Gene Ontology Biological Process terms across neuronal gene modules associated with fetal and adult mDA neurons. Grey nodes represent enriched GO terms, with node outline color indicating whether the term is supported by adult-associated modules (red) or fetal-associated modules (purple). Coloured nodes represent genes contributing to the enriched terms and are shaded according to whether they derive from adult-enriched or fetal-enriched modules. Edges connect GO terms to overlapping module genes. Only significantly enriched neuronal-related GO terms passing the FDR threshold of 0.001 are shown. **e,** Correlation of differentiation protocols to fetal and adult dopaminergic reference states. Each point represents the average mDA profile for a timepoint–protocol–publication (TPP), positioned according to its correlation with fetal (x-axis) and adult (y-axis) *in vivo* reference signatures. Gene module scores were first corrected for single-cell versus single-nuclei technology bias, and average module profiles were then computed for each TPP before calculating correlations with the fetal and adult references. Star symbols indicate the fetal and adult reference anchors, corresponding to the expected positions of a perfect fetal or adult reference relative to the fetal–adult correlation. Only protocols with at least 10 mDA neurons were plotted. **f,** Boxplot plots showing per-cell distributions and mean scores for glycolysis, hypoxia, ER stress, mDA fidelity, and maturation across timepoint–protocol–publication. Each point represents timepoint–protocol–publication (TPP) for each model. Scores were calculated using gene-set module scoring and corrected for single-cell versus single-nuclei technology effects. The mDA fidelity score was derived from top 50 marker genes for fetal MB mDA neurons, and the maturation score was based on Module 8 maturation-associated genes and adult-enriched TFs. **g,** Scatter plots showing relationships between glycolysis, hypoxia, and ER stress scores and mDA identity or maturation across TPPs, after removing between-protocol/publication effects and controlling for culture time using a second-degree polynomial regression. Black lines indicate fitted linear trends. Insets show Spearman correlation coefficients, *P* values, and the number of TPPs included in each comparison.

